# Distance-based Reconstruction of Protein Quaternary Structures from Inter-Chain Contacts

**DOI:** 10.1101/2021.05.24.445503

**Authors:** Elham Soltanikazemi, Farhan Quadir, Raj S. Roy, Jianlin Cheng

## Abstract

Predicting the quaternary structure of a protein complex is an important and challenging problem. Inter-chain residue-residue contact prediction can provide useful information to guide the *ab initio* reconstruction of quaternary structures of protein complexes. However, few methods have been developed to build quaternary structures from predicted inter-chain contacts. Here, we introduce a new gradient descent optimization algorithm (GD) to build quaternary structures of protein dimers utilizing inter-chain contacts as distance restraints. We evaluate GD on several datasets of homodimers and heterodimers using true or predicted contacts. GD consistently performs better than a simulated annealing method and a Markov Chain Monte Carlo simulation method. Using true inter-chain contacts as input, GD can reconstruct high-quality structural models for homodimers and heterodimers with average TM-score ranging from 0.92 to 0.99 and average interface root mean square distance (I-RMSD) from 0.72 Å to 1.64 Å. On a dataset of 115 homodimers, using predicted inter-chain contacts as input, the average TM-score of the structural models built by GD is 0.76. For 46% of the homodimers, high-quality structural models with TM-score >= 0.9 are reconstructed from predicted contacts. There is a strong correlation between the quality of the reconstructed models and the precision and recall of predicted contacts. If the precision or recall of predicted contacts is >20%, GD can reconstruct good models for most homodimers, indicating only a moderate precision or recall of inter-chain contact prediction is needed to build good structural models for most homodimers. Moreover, the accuracy of reconstructed models positively correlates with the contact density in dimers and depends on the initial model and the probability threshold of selecting predicted contacts for the distance-based structure optimization.

## Introduction

Determination of interactions between protein chains in a protein complex is important for understanding protein function and cellular processes and can play significant roles in designing and discovering new drugs^1^. Detailed protein-protein interactions are represented by the three-dimensional shape of a complex consisting of interacting proteins (i.e., quaternary structure). Experimental techniques such as X-ray crystallography and nuclear magnetic resonance (NMR) can determine the quaternary structure of protein complexes with high accuracy. However, these experimental approaches are costly and time-consuming, and therefore cannot be applied to most protein complexes. Therefore, computational modeling approaches, which provide a faster and inexpensive way to predict quaternary structures, have become increasingly popular and important^2^.

Computational protein docking, currently the most widely used approach for modeling complex structures, takes the tertiary structures of individual proteins as input to build the quaternary structure of the complex as output^3-9^. Docking methods can be largely divided into two categories including template-based modeling, in which known protein complex structures in the Protein Data Bank (PDB) are used as templates^10-17^ to guide modeling, and template-free modeling (*ab initio* docking), which does not use any known structure as template, and instead searches through a large conformation space for relative orientations of protein chains with minimum binding energy. The binding energy is often roughly approximated by geometric and electrostatic complementarity, inter-chain hydrogen binding, hydrophobic interactions, and residue-residue contact potentials^18-23^.

Although template-based docking works well if a good structural template is available, it cannot be applied to most protein complexes that lack suitable templates^2,24^. *Ab initio* docking methods can predict the quaternary structure of acceptable quality for some protein complexes, but according to several rounds of Critical Assessments of Predictions of Interactions (CAPRI), they still cannot achieve adequate accuracy for most protein complexes^24,25^. One main reason for the low accuracy is that the *ab initio* docking methods need to search through a huge conformation space, which is usually not feasible with limited time and computing resources. To reduce the search space, several methods started to use the interface contacts between proteins to constrain conformation search^26-30^ and were able to enhance docking accuracy^30^, showing inter-chain (inter-protein) contacts can provide valuable information to build protein quaternary structures as what had happened in protein tertiary structure prediction.

The major advances of *ab initio* tertiary structure prediction of a single protein chain have been largely driven by accurate prediction of intra-chain residue-residue contact prediction and the development of methods of reconstructing tertiary structures from the contacts^31-36^. However, there are still very few methods available to reconstruct protein quaternary structures from predicted inter-chain residue-residue contacts. With the emergency of inter-chain contact prediction enhanced by residue-residue co-evolutionary analysis and deep learning^37-41^, it is crucial to create robust methods to efficiently and effectively use inter-chain contacts to directly reconstruct protein quaternary structures.

Gradient descent optimization has become a popular method to build the tertiary structure of proteins using intra-protein (intra-chain) residue-residue contacts or distances. AlphaFold^31^, which was ranked first in CASP13, developed a gradient descent-based folding method to generate protein tertiary structure from intra-chain distances. trRosetta^32^, a powerful tool for protein tertiary structure modeling, uses a gradient descent-like method (MinMover from pyRosetta) to build the structure of individual proteins from predicted residue-residue distances. A recent protein folding framework based on gradient descent, GDFOLD^42^, uses intra-chain contacts as input constraints to directly optimize the positions of C_α_atoms of a protein.

Motivated by the recent success of applying gradient descent to protein tertiary structure prediction, in this study, we develop an *ab initio* gradient descent optimization method (GD) to construct quaternary structures of protein dimers from inter-chain contacts. We first test if the proposed method can generate high quality structures of protein dimers using true contacts. Then, we apply it to construct quaternary structures of homodimers from predicted, noisy, and incomplete contacts. To rigorously benchmark its performance, we also implement a Markov Chain simulation method (MC) based on RosettaDock^43^ and apply a simulated annealing method based on Crystallography and NMR System (CNS)^41^ to reconstruct protein complex structures from inter-protein contacts and compare them with GD. We evaluate the three methods on several in-house datasets consisting of 233 homodimers and heterodimers in total as well as on a standard dataset of 32 heterodimers^40,44^ with true or predicted contacts. GD consistently performs better than MC and CNS on all the datasets. It is able to reconstruct high-quality structures from true inter-chain contacts and good structures for most homodimers when predicted contacts are only moderately accurate.

## Results and Discussions

### Reconstruction of quaternary structure from native (true) contacts

We first apply GD, MC and CNS to generate quaternary structures for 44 homodimers in the Homo44 dataset using true inter-chain contacts as input. The models reconstructed by the methods are evaluated by five complementary metrics against known experimental structures of the homodimers: root-mean-square deviation (RMSD), TM-score, the percentage of native contacts existing in predicted models (f_nat), interface RMSD (I_RMSD), and Ligand RMSD (L_RMSD) widely used in the field.

The detailed results of GD on the Homo44 dataset in terms of TM-score, RMSD, f_nat, I_RMSD, and L_RMSD) and the length and number of contacts of the homodimers are reported in supplemental **Table S1**. GD is able to generate high-quality structural models for all the dimers when true inter-chain contacts are provided as input. For instance, TM-score of the models ranges from 0.936 to 0.999 and I_RMSD from 0.204 Å to 1.85 Å. The average of RMSD, TM-score, f_nat, I_RMSD, and L_RMSD of GD, MC and CNS is compared in **Table 1** (see the per-dimer comparison of the three methods in terms of each metric in supplemental **Figures S1-S5**). GD performs best in terms of all the metrics, while MC performs better than CNS. The average RMSD of GD is 0.63Å, which is lower than 0.76Å of MC and 1.16Å of CNS. The average TM-score of GD is 0.99 – an almost perfect score, which is higher than 0.98 of MC and 0.91 of CNS. Moreover, GD realizes 92.19% of native contacts (f_nat = 92.19%), higher than 91.39% of MC and 82.49% of CNS. The average I_RMSD and L_RMSD of GD are 0.77Å and 1.38Å, lower than those of the other two methods. **Figure 1** illustrates high-quality structural models reconstructed by GD, MC, and CNS that are superimposed with the true structure of a dimer (PDB code: 1XDI) in Homo44.

**Table 1.** Mean and standard deviation (std) of RMSD, TM-score, f_nat, I_RMSD, and L_RMSD of the three methods on 44 homodimers in Homo44.

| Evaluation Metric | GD | MC | CNS |
| --- | --- | --- | --- |
| **RMSD (mean, std)** | **0.63<sup>+0.3788</sup>** | 0.76<sup>+0.361</sup> | 1.16<sup>+1.0043</sup> |
| **TM-score (mean, std)** | **0.99<sup>+0.0132</sup>** | 0.98<sup>+0.014</sup> | 0.91<sup>+0.0102</sup> |
| **f_nat (mean, std)** | **92.19<sup>+8.64</sup>** | 91.39<sup>+9.08</sup> | 82.49<sup>+22.02</sup> |
| **I_RMSD (mean, std)** | **0.77<sup>+1.05</sup>** | 1.35<sup>+3.98</sup> | 12.46<sup>+8.46</sup> |
| **L_RMSD (mean, std)** | **1.38<sup>+0.8</sup>** | 1.7<sup>+0.9</sup> | 11.18<sup>+14.51</sup> |

**Figure 1.**
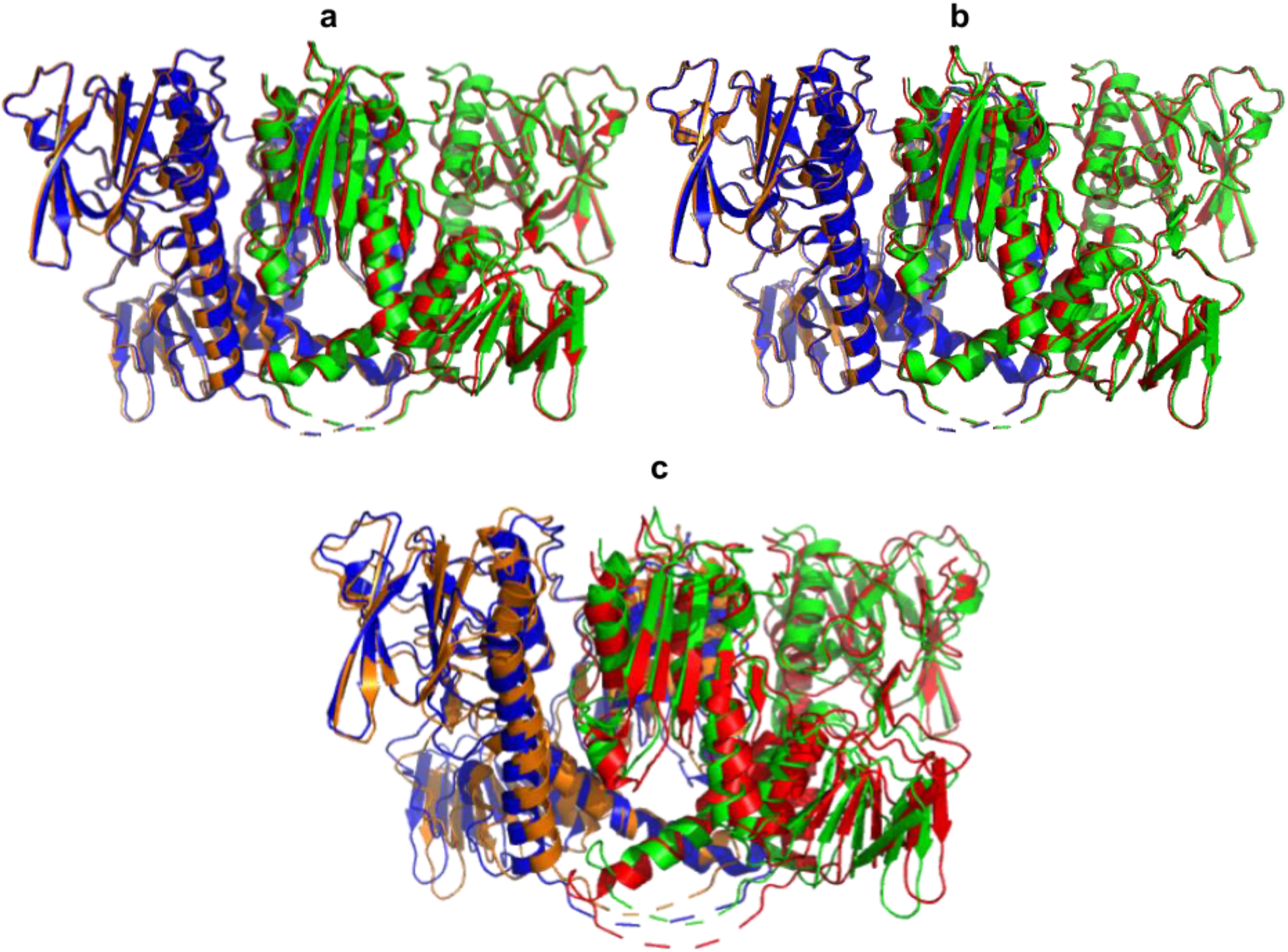
The superposition of the native structure of 1XDI and the models reconstructed by three methods (i.e., green and orange denoting the true dimer structure and blue and red the reconstructed dimer structure): (a) GD, (b) MC and (c) CNS. TM-score, RMSD, f_nat, I_RMSD, L_RMSD of the model predicted by GD are 0.99, 0.56Å, 94.52%, 0.24Å, and 0.74Å, respectively. TM-score, RMSD, f_nat, I_RMSD, L_RMSD of the model predicted by MC are 0.99, 0.61Å, 93.15%, 0.45Å, and 1.29Å, respectively. TM-score, RMSD, f_nat, I_RMSD, L_RMSD of the model predicted by CNS are 0.88, 2.25Å, 74.79%, 1.49Å, and 5.18Å, respectively.

We also evaluate GD with MC and CNS on 73 heterodimers in the Hetero73 dataset using true inter-chain contacts as input. The detailed per-dimer results of GD are shown in supplemental **Table S2**. A comparison of the three methods is shown in **Table 2**. The average RMSD, I_RMSD, and L_RMSD of GD are lower than the other two methods, while its average TM_score and f_nat are higher than the other two methods, indicating that GD performs best, while MC works better than CNS. A per-dimer comparison of RMSD and TM-score of the models reconstructed by the three methods is depicted in **Figures S6** and **S7**, respectively. The models reconstructed by GD for the heterodimers have high quality on average (e.g., mean RMSD = 1.23Å and TM-score = 0.92). However, in comparison with the results on homodimers in **Table 1**, the average accuracy on heterodimers is lower than that on homodimers. A main reason is that heterodimers tend to have lower inter-chain contact density (i.e., # of inter-chain contacts / sum of the sequence lengths of two chains in a dimer)^41,45^ on average, leading to fewer distance restraints available for structure reconstruction.

**Table 2.** Mean and standard deviation (std) of RMSD, TM-score, f_nat, I_RMSD, and L_RMSD results of the three methods on 73 heterodimers in the Hetero73 dataset.

| Evaluation Metric | GD | MC | CNS |
| --- | --- | --- | --- |
| **RMSD (mean, std)** | **1.23<sup>+1.91</sup>** | 4.76<sup>+8.01</sup> | 7.7<sup>+12.99</sup> |
| **TM-score (mean, std)** | **0.92<sup>+0.12</sup>** | 0.85<sup>+0.16</sup> | 0.79<sup>+0.23</sup> |
| **F_nat (mean, std)** | **90.31<sup>+16.77</sup>** | 82.59<sup>+26.68</sup> | 84.43<sup>+23</sup> |
| **I_RMSD (mean, std)** | **0.72<sup>+1.02</sup>** | 1.58<sup>+1.7</sup> | 1.65<sup>+4.51</sup> |
| **L_RMSD (mean, std)** | **3.75<sup>+6.15</sup>** | 7.78<sup>+11.8</sup> | 9.21<sup>+14.05</sup> |

Moreover, we evaluate the three methods on 32 heterodimers in the Std32 dataset. The detailed results of GD are presented in supplemental **Table S3**. The average TM-score, RMSD, f_nat, I_RMSD, and L_RMSD of the models reconstructed by GD, MC, and CNS are reported in **Table**

**3**. Similar to the results on the other datasets, GD generates high-quality models on average and performs best in terms of all the metrics, while MC performs substantially better than CNS.

### Analysis of two key factors impacting the quality of models reconstructed by GD from native contacts

We have observed that the quality of the generated structures is affected by two factors: initial structure in the optimization and inter-chain contact density in a dimer.

**Figure 2** shows the changes of TM-score and RMSD of 20 models simulated by GD with different start models for a dimer 1Z3A in Homo44. The TM-score of the final models ranges from about 0.55 to about 1.0, depending on the start models. Given a reasonable initial structure, GD converges to a high-quality local minima that has similar performance as the global minima, generating an accurate structure with an almost perfect TM-score = 1. On the other hand, starting from a poor initial model, the algorithm can get stuck in a bad local minima, producing a low-quality model. Therefore, it is useful to run GD multiple times with different start models. Based on the experiment on Homo44 and Hetero73 datasets, using 20 different start models to run GD 20 times is able to build almost perfect quaternary structural models with TM-score = 0.99 and an RMSD less than 1Å from true inter-chain contacts for most dimers (see **Table S1** and **Table S2** for details).

**Figure 2.**
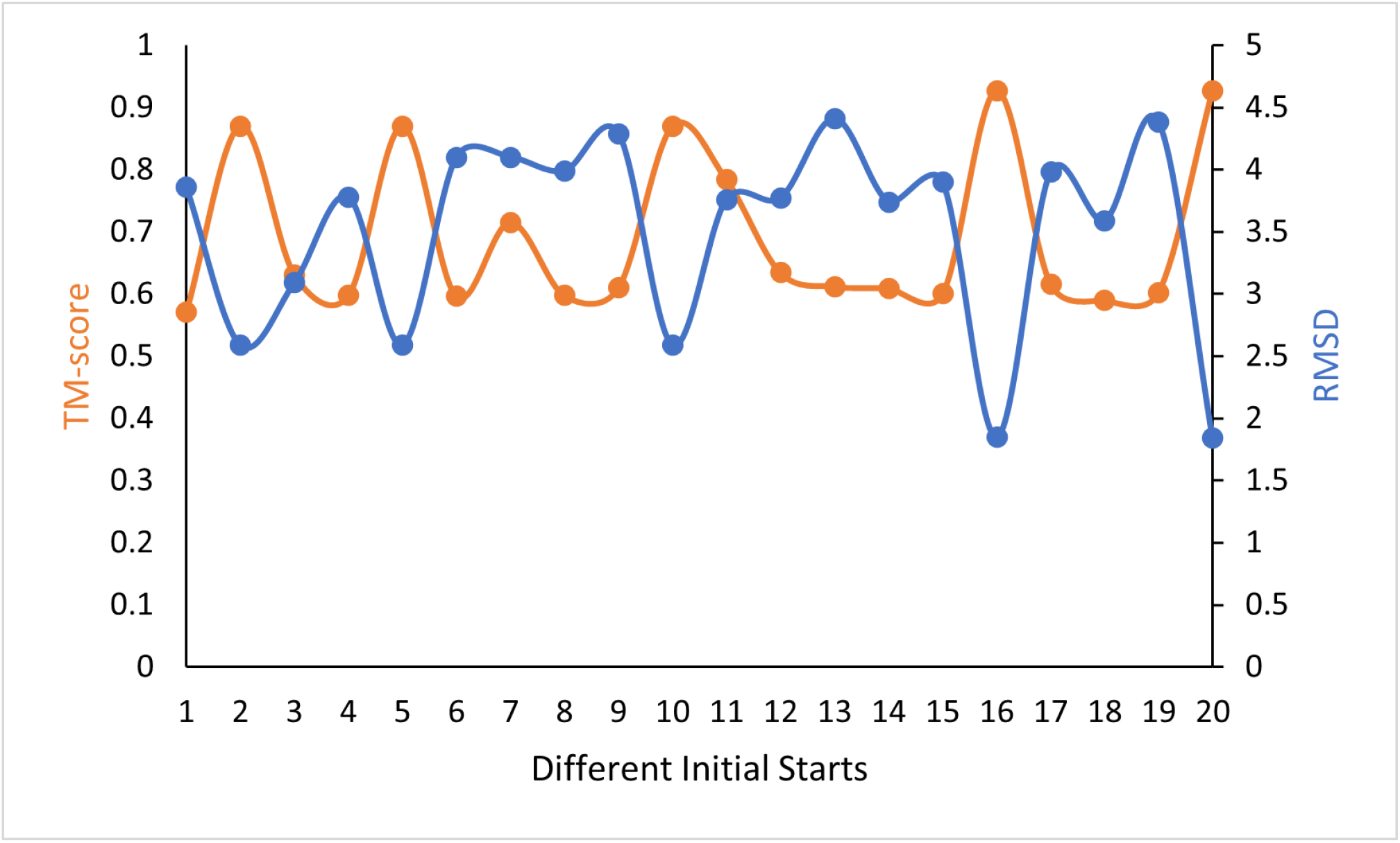
TM-score (orange dots) and RMSD (blue dots) of 20 models reconstructed from 20 different initial structures by GD for a homodimer 1Z3A. X-axis denotes the indices of 20 models constructed with different start structures. Y-axis denotes TM-score or RMSD of the models.

In addition to initial models, the contact density of a dimer strongly influences the quality of the models reconstructed from native contacts. **Figure 3** illustrates how TM-score and RMSD of the models reconstructed for 73 heterodimers change with respect to the density of true contacts. When the contact density is above ∼0.25, almost all the models have a very low RMSD (< 1Å) and a very high TM-score (close to 1). When contact density is lower than ∼0.25, there are both good-quality and low-quality models. Overall, with an increase of the contact density, the quality of the reconstructed structure increases in terms of all the metrics: RMSD, TM-score, f_nat, I_RMSD, and L_RMSD (results for f_nat, I_RMSD, and L_RMSD not shown).

**Figure 3.**
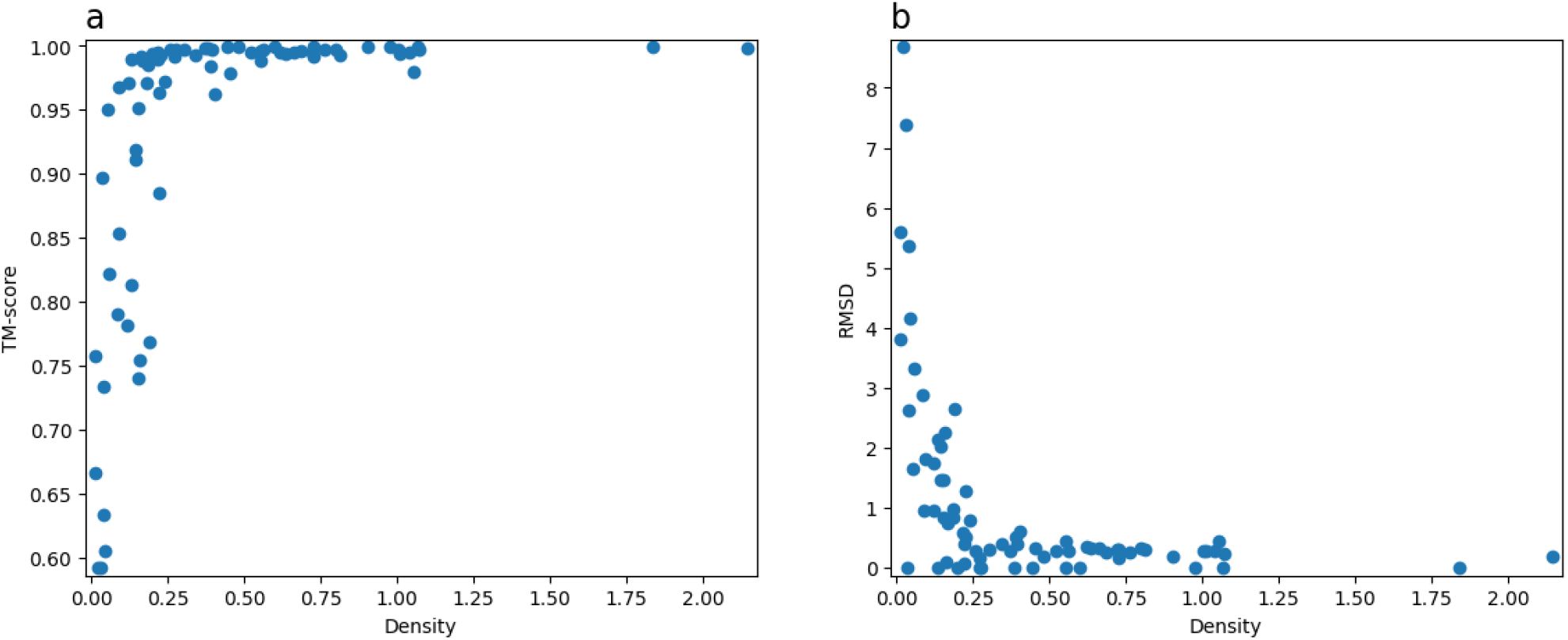
TM-scores and RMSD of the models versus the inter-chain contact density of 73 heterodimers.

### Reconstruction of quaternary structures of homodimers from predicted inter-chain contacts

We evaluate the performance of the three optimization methods on homodimers using predicted inter-chain contacts because the newly developed deep learning methods such as ResCon can make inter-chain contact prediction with reasonable accuracy for a large portion of homodimers. The three methods are compared on three subsets (Set A, Set B, Set C) of homodimers in the Homo115 dataset. Set A consists of 40 dimers with small interaction interfaces. Set B has 37 dimers with medium interaction interfaces. Set C contains 38 complexes with large interaction interfaces. The detailed results of GD (average TM-score, RMSD, f_nat, I_RMSD, and L_RMSD) as well as the precision and recall of the predicted contacts for sets A, B, and C are shown in supplemental **Tables S4, S5** and **S6**, respectively. The precision of predicted inter-chain contacts is measured by 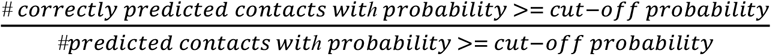, and the recall of predicted inter-chain contacts by 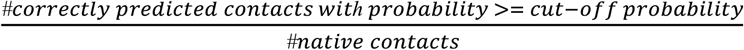, where *cut* – *off probability* of selected contacts is set to 0.5.

The average performance of GD, MC, and CNS on Sets A, B, and C is compared in **Table 4, Table 5**, and **Table 6**, respectively. Similar as observed on models reconstructed from true inter-chain contacts, GD performs best here, MC second, and CNS third in terms of almost all the evaluation metrics. Moreover, the average accuracy generally increases with the increase of the size of the interaction interfaces (i.e., accuracy of Set C > accuracy of Set B > accuracy of Set A), showing that it is easier to reconstruct quaternary structures with larger interaction interfaces. The average TM-score of the structural models built for the three datasets by GD is 0.68, 0.80, and 0.81, respectively, higher than the models predicted by MC and CNS. GD generates models with higher TM-score for most dimers. The average TM-score of the models reconstructed by GD for all 115 homodimers in Set A, Set B, and Set C is 0.76. Moreover, for 53 out of 115 (46%) homodimers, the models reconstructed by GD have high TM-scores (>= 0.9) (see Tables S4, S5 and S6), suggesting that GD is able to reconstruct high-quality models for a large portion of dimers using only predicted inter-chain contacts as input. **Figure 4** illustrates a high-quality model reconstructed for dimer 1C6X (precision of contact prediction = 40.24% and recall of contact prediction = 49.28%, TM-score = 0.99, f_nat = 84.61%).

**Table 3.**
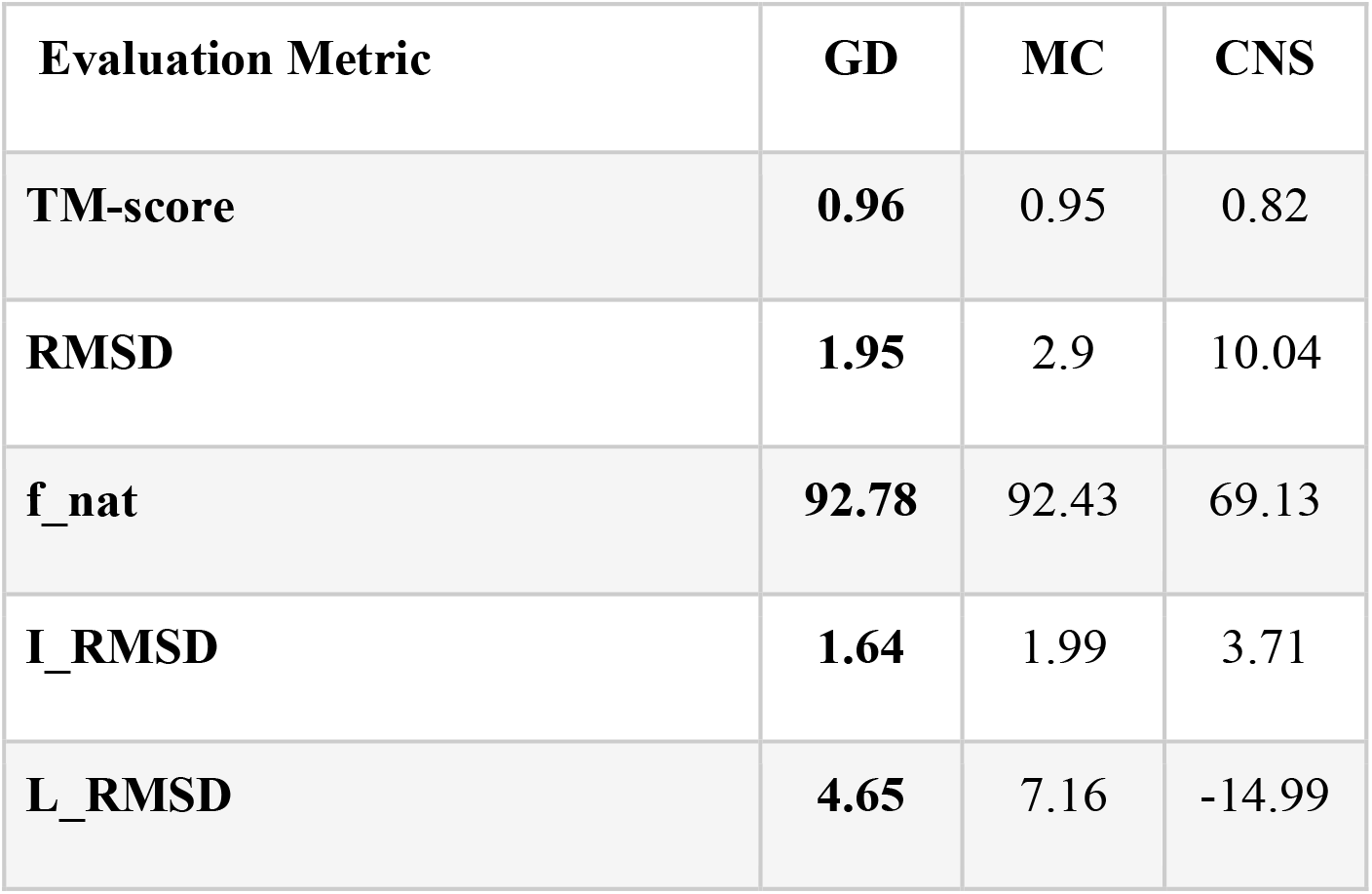
Average RMSD, TM-score, f_nat, I_RMSD, and L_RMSD of GD, MC and CNS on 32 dimers in the Std32 dataset.

| Evaluation Metric | GD | MC | CNS |
| --- | --- | --- | --- |
| **TM-score** | **0.96** | 0.95 | 0.82 |
| **RMSD** | **1.95** | 2.9 | 10.04 |
| **f_nat** | **92.78** | 92.43 | 69.13 |
| **I_RMSD** | **1.64** | 1.99 | 3.71 |
| **L_RMSD** | **4.65** | 7.16 | -14.99 |

**Table 4.**
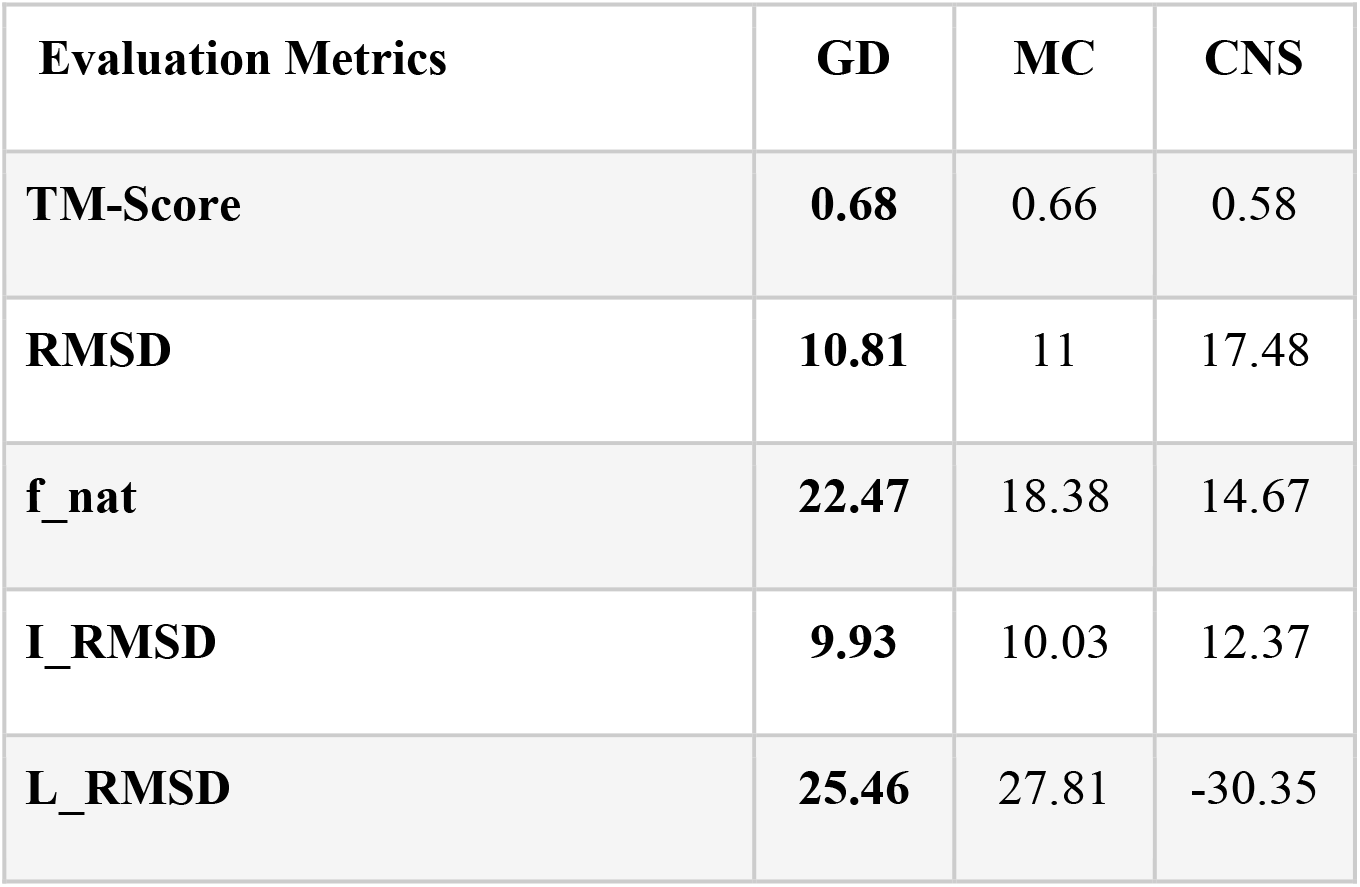
Average RMSD, TM-score, f_nat, I_RMSD, and L_RMSD of the best models reconstructed by the three methods for the homodimers in Set A using predicted contacts as input.

**Table 5.**
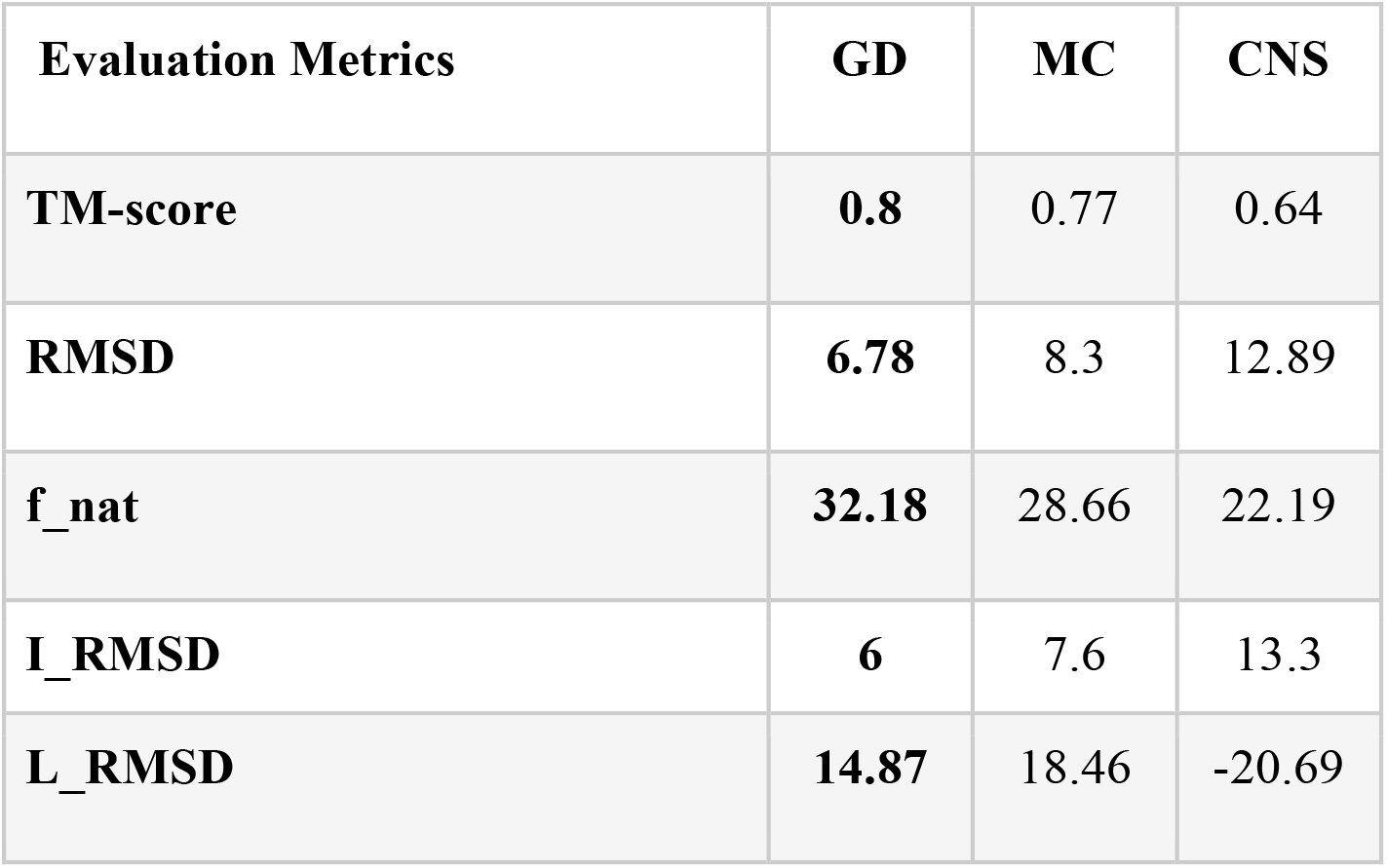
Average RMSD, TM-score, f_nat, I_RMSD, and L_RMSD of the best models reconstructed by the three methods for Set B with predicted inter-chain contacts as input.

**Table 6.** Average RMSD, TM-score, f_nat, I_RMSD, and L_RMSD of the best models reconstructed by the three methods for Set C with predicted contacts as input.

| **Evaluation Metrics** | **GD** | **MC** | **CNS** |
| --- | --- | --- | --- |
| **TM-score** | **0.81** | 0.80 | 0.76 |
| **RMSD** | **6.26** | 6.77 | 9.5 |
| **f_nat** | 37.43 | 35.07 | **42.3** |
| **I_RMSD** | **5.01** | 5.46 | 7.41 |
| **L_RMSD** | **12.73** | 13.96 | -16.3 |

**Figure 4.**
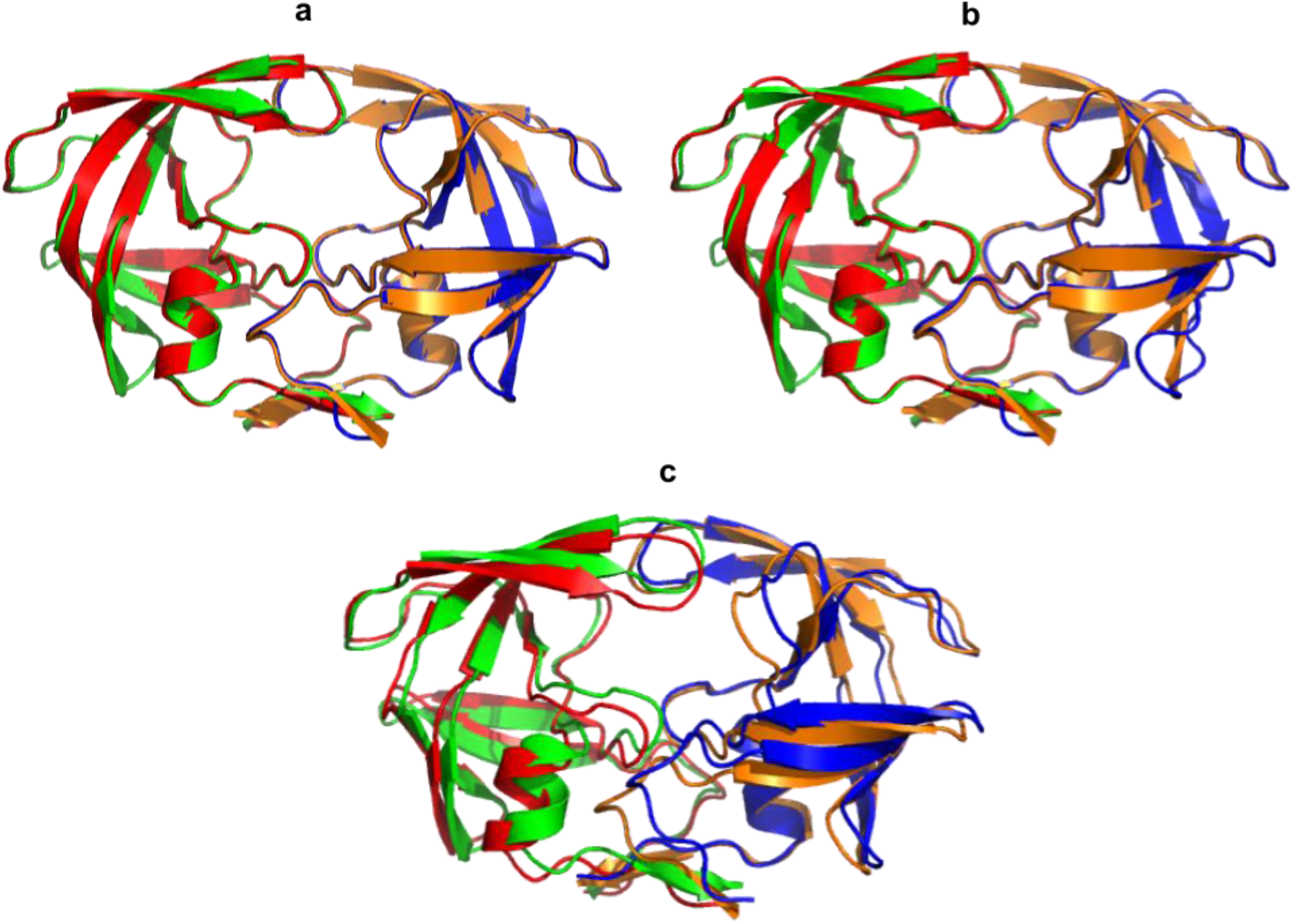
The superposition of the native structure of 1C6X and the models generated by three methods (i.e., green and orange representing the true dimer structure and blue and red the generated models): (a) GD, (b) MC and (c) CNS. TM-score, RMSD, f_nat, I_RMSD, L_RMSD of the model predicted by GD are 0.99, 0.4Å, 84.61%, 0.4Å, and 0.91Å, respectively. TM-score, RMSD, f_nat, I_RMSD, L_RMSD of the model predicted by MC are 0.98, 0.6Å, 78.84%, 0.6Å, and 1.6Å, respectively. TM-score, RMSD, f_nat, I_RMSD, L_RMSD of the model predicted by CNS are 0.86, 2.02Å, 41.6%, 2.14Å, and 5.68Å, respectively.

We investigate the relationship between the quality of the models generated by GD and the precision and recall of predicted contacts. **Figure 5a** plots the TM-score of the models constructed for the dimers in Homo115 against the precision of contacts predicted for them. The correlation between the two is 0.78, indicating that the quality of the structural models increases with respect to the precision of predicted contacts. It is worth noting that if the precision is > 20%, most reconstructed models have good quality (e.g., with TM-score > 0.8 or even close to 1). If the precision is > 40%, all the models have good quality (TM-score > 0.8). The results demonstrate that there is no need to get a very high accuracy of contact prediction for GD to obtain high-quality structural models for homodimers as long as its accuracy reaches a specific threshold. GD is robust against the noise in predicted contacts. This result is encouraging news for the community to develop more methods to predict inter-chain contacts in protein complexes.

**Figure 5.**
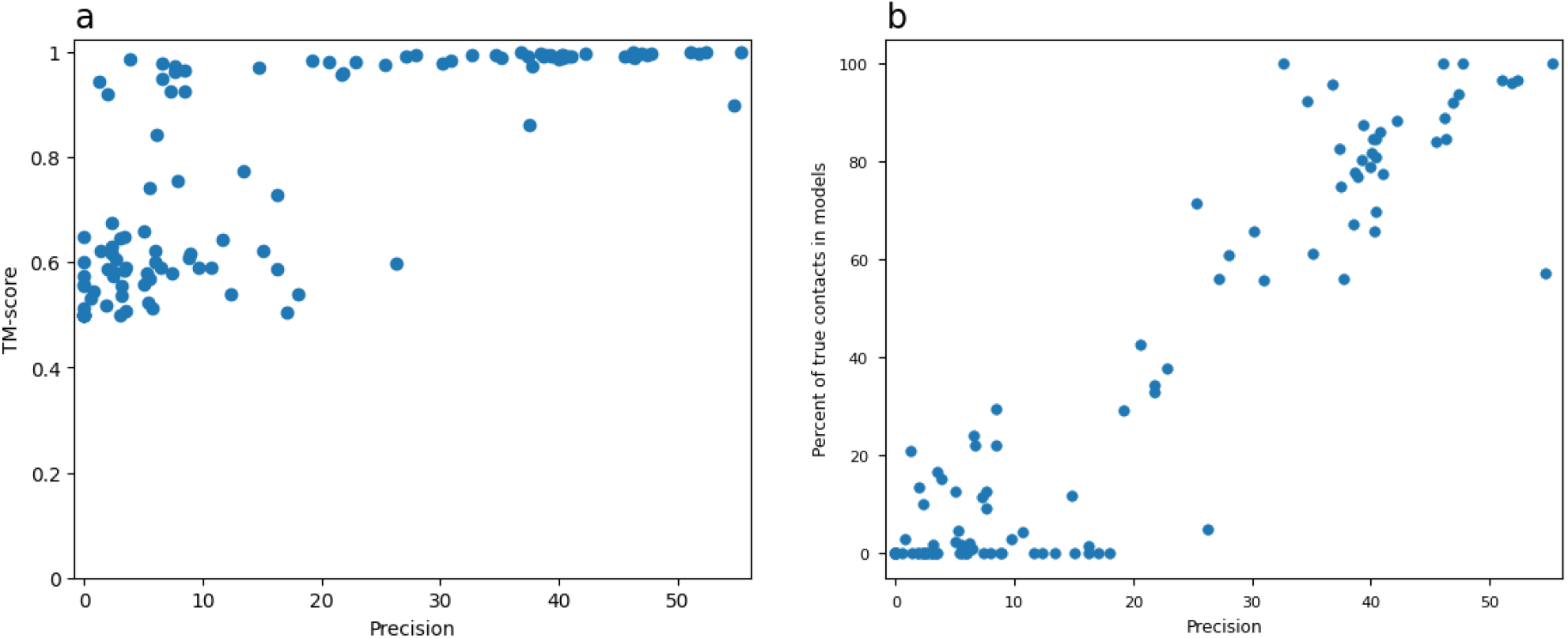
The plot of TM-score and percent of native contacts of the models (f_nat) against the precision of predicted contacts on Homo115 dataset. Pearson’s correlation between the two is 0.78.

**Figure 5b** also reveals the strong positive correlation between the percent of true contacts existing in the reconstructed structural models and the precision of predicted contacts. Pearson’s correlation between the two is 0.94. Moreover, when the precision of predicted contacts is > 40%, a high percent (>50%) of native contacts are realized in the models.

Furthermore, there is a strong correlation between the quality of reconstructed models (e.g., TM-Score and f_nat) and the recall of the predicted inter-chain contacts as shown in **Figure 6**. Pearson’s correlation between TM-score and recall is 0.78 and between f_nat and recall is 0.93, showing that a higher recall of predicted contacts leads to better reconstructed models. As shown in **Figure 6a**, when the recall of predicted contacts is >20%, all the reconstructed models except a few cases have good quality, i.e., their TM-score is > 0.8 and even close to 1, indicating only a small portion of true contacts are needed to build good quaternary structural models for most homodimers. Even if the recall of predicted contacts is < 20%, good models (TM-score > 0.8) can still be reconstructed for some dimers. Moreover, as shown in **Figure 6b**, when the recall of predicted contacts is >20%, the percent of true contacts (f_nat) in the models reconstructed for all but a few dimers is higher than the recall of predicted contacts that are used as input, indicating that the optimization process of GD can realize (recall) more true contacts than what is provided in the predicted input contacts.

**Figure 6.**
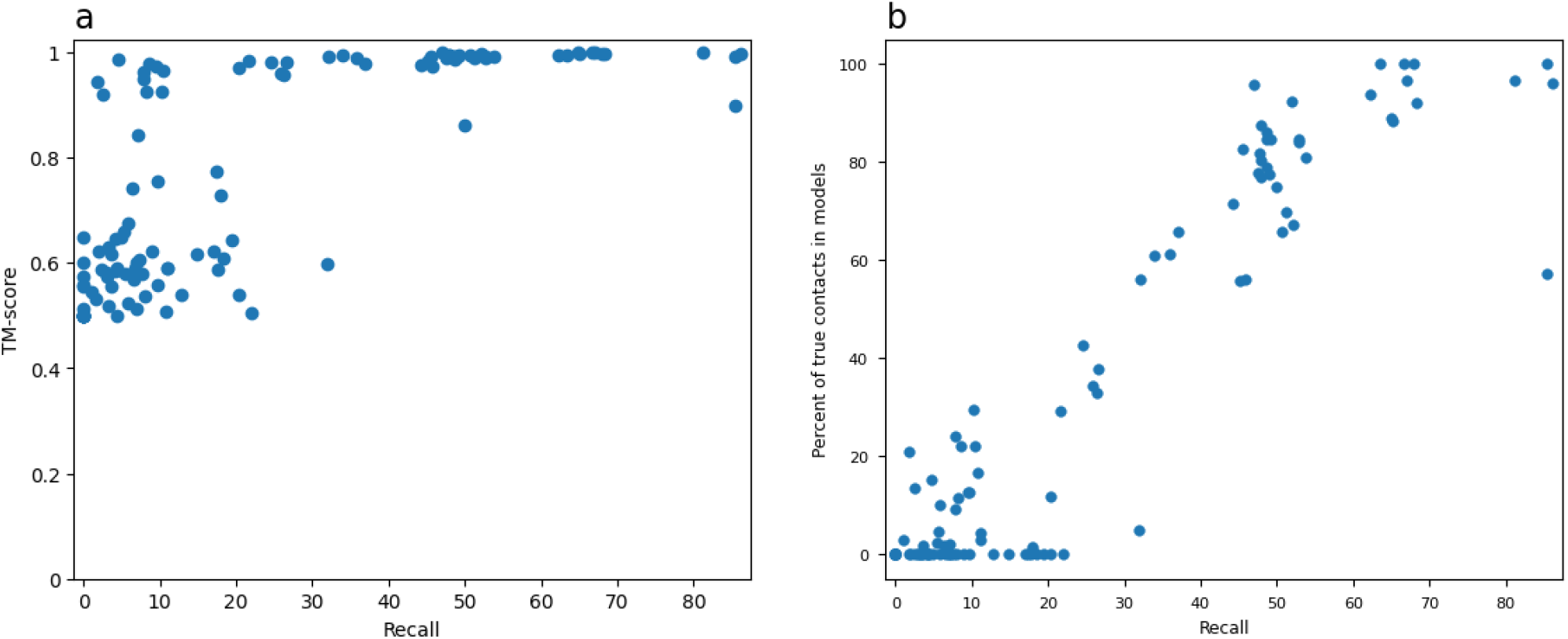
TM-score and percent of native contacts of the predicted models (f_nat) reconstructed by GD versus the recall of the predicted inter-chain contacts on the Homo115 dataset.

Moreover, we investigate how the cut-off probability of selecting predicted inter-chain contacts as input affects the quality of reconstructed structural models. To determine good cut-off probabilities for selecting predicted contacts, we test different cut-off values in the range [0.3, 0.9], with a step size of 0.1. **Figure 7** shows how the average TM-score and RMSD of reconstructed models change with respect to the cut-off probabilities on the Homo115 dataset. The best model quality (lowest RMSD and highest TM-score) is reached at the cut-off probability of 0.5 on the dataset. We imagine that the best cut-off probability can be data- and method-dependent. Therefore, it can be useful to try different cut-off probabilities to reconstruct models and then select good ones from them on different datasets.

**Figure 7.**
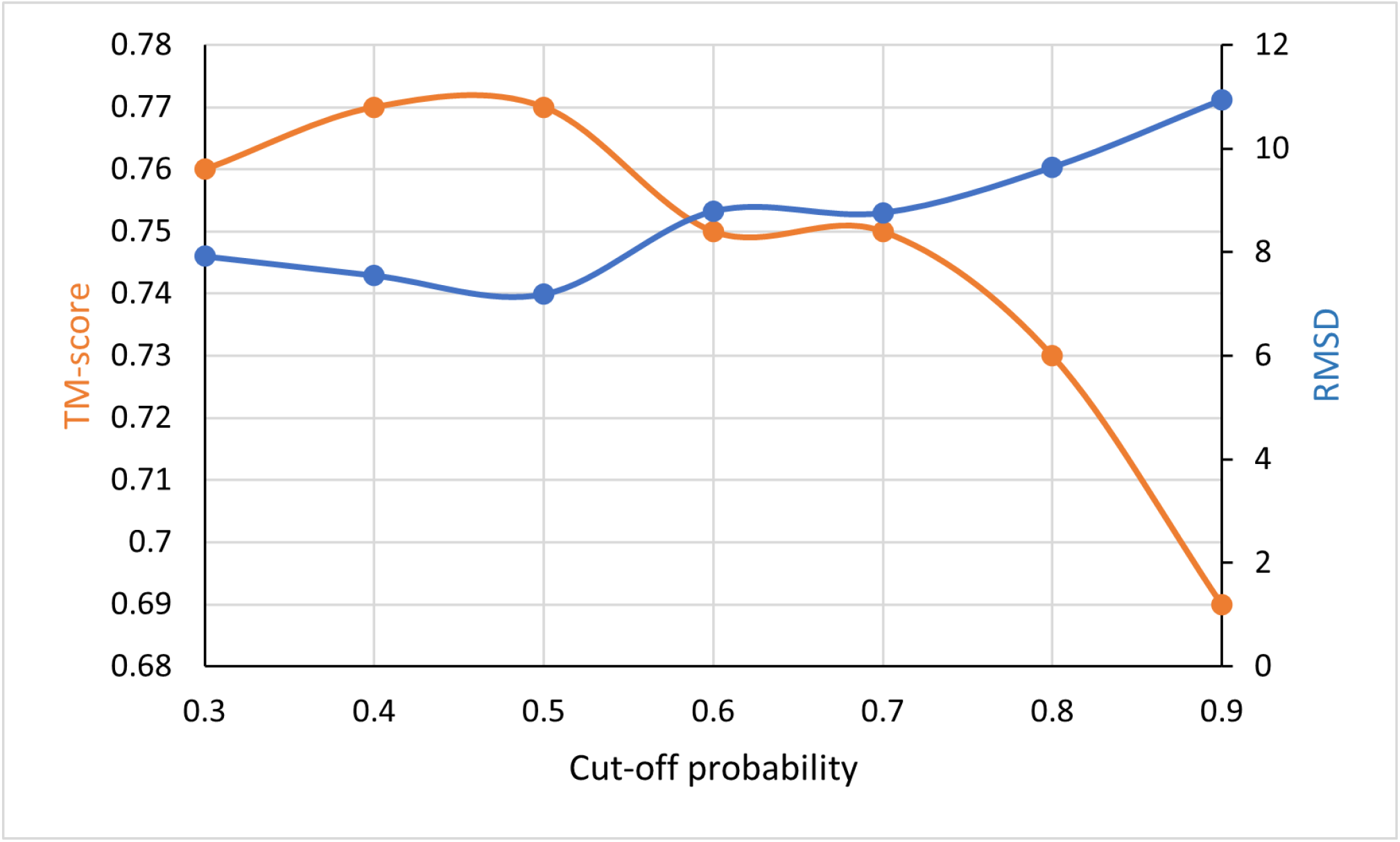
The average RMSD and TM-score of models reconstructed for homodimers in the Homo115 dataset versus the cut-off probability of selecting predicted inter-chain contacts as restraints.

## Conclusion

We design and develop a gradient descent distance-based optimization (GD) method to reconstruct quaternary structure of protein dimers from inter-protein contacts and compare it with the Markov Chain Monte Carlo and simulated annealing optimization methods adapted to address the problem. GD performs consistently better than the other two methods in reconstructing quaternary structures of dimers from either true or predicted inter-chain contacts. GD can reconstruct high-quality structures for almost all homodimers and heterodimers from true inter-chain contacts and can build good structural models for many homodimers from only predicted inter-chain contacts, demonstrating distance-based optimizations are useful tools for predicting the quaternary structures. Moreover, we show that the contact density, size of interaction interface, precision and recall of predicted contacts, and threshold of selecting contacts as restraints influence the accuracy of reconstructed models. Particularly, when the precision and recall of predicted contacts reach a moderate level (e.g., >20%), GD can construct good models for most homodimers, demonstrating that predicting inter-chain contacts (or even distances) and distance-based optimization are a promising *ab initio* approach to predicting the quaternary structures of protein complexes.

## Materials and Methods

### Inter-Chain Contacts and Dimer Datasets

Two residues from two protein chains in a dimer are considered an inter-chain contact if any two heavy atoms from the two residues have a distance less than or equal to 6 Å^41,45^. True contacts of a dimer with the known quaternary structure in the PDB are identified according to the coordinates of atoms in the PDB file of the dimer.

We use several in-house datasets of protein homodimers and heterodimers with true and/or predicted inter-protein contacts as well as a standard dataset consisting of 32 heterodimers (Std32)^46^ in this study. The first in-house dataset has 44 homodimers randomly selected from the Homo_Std^41^ curated from the 3DComplex database^47^, each of which have 39 to 621 true contacts (called Homo44). The second in-house data includes 115 homodimers (called Homo115) selected from Homo_Std, each of which has at least 21 predicted inter-chain contacts with a probability of >=0.5. Our in-house deep learning method - ResCon^46^ is applied to predict inter-chain contacts for the dimers in Homo115. Homo115 is divided into three subsets (Set A, Set B, and Set C) according to the size of interfaces. Set A has 40 protein complexes with small interaction interfaces consisting of 14 to 68 true inter-chain contacts. Set B consists of 37 complexes having medium interaction interfaces with 69 to 129 true contacts. Set C consists of 38 complexes having large interaction interfaces with 131 to 280 true contacts. The third in-house dataset contains 73 heterodimers (called Hetero73)^46^ curated from the PDB, in which the sum of the lengths of the two chains is less than or equal to 400. The heterodimers in Hetero73 have 2 to 255 true inter-protein contacts.

### Gradient Descent Cost Function and Optimization

The inter-chain contacts are used as distance restraints for the gradient descent method to build the structures of protein dimers. The cost function to measure the satisfaction of the distance between any two residues in contact to guide the structural modeling is defined as follows:

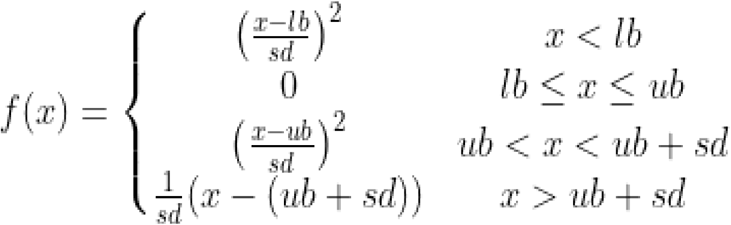

Here, *lb* and *ub* represent the lower bound and upper bound of the distance (*x*) between two residues that are assumed to be in contact. As mentioned earlier, two residues are considered to be in contact if the distance between their heavy atoms is less than 6 Å. However, to simplify the process of restraint preparation, two residues are considered to be in contact if the distance between their *C*_*β*_ atoms (*C*_*α*_ for Glycine) is less than 6 Angstrom. The lower bound (*lb*) is empirically set to 0 and the upper bound (*ub*) to 6 Å. *sd* is the standard deviation, which is set to 0.1. Based on this cost function, if the distance between two residues in contact is <= 6 Å, i.e., the contact restraint is satisfied, and the cost is 0.

The complete contact cost function for a structural model of a dimer to be minimized is the sum of the costs for all contacts used in modeling (called contact energy). For simplicity, all restraints have equal weights and play equally important roles in modeling. The contact energy function is differentiable with respect to the distances between residues and coordinates of atoms of the residues, and therefore it can be minimized by a gradient descent iterative algorithm (GD), i.e., Limited-memory Broyden–Fletcher–Goldfarb–Shanno algorithm (L-BFGS)^31,48^) used in this study.

We implement GD using pyRosetta. The total energy function for the structural optimization is the combination of the contact energy and the talaris2013 potentials^31^. The input to the algorithm includes inter-chain contacts and an initial random conformation of a dimer. An initial conformation of a protein dimer is generated by making 40 random rotations and translations ranging from *1*° – *360*° and *1 Å* – *20 Å*, respectively. Specifically, the tertiary structure of each protein chain is rotated and translated arbitrarily along the line connecting the centers of the two chains, aiming to make the two protein chains facing each other.)

Then 6000 iterations of the gradient descent optimization (i.e., L-BFGS) are carried out to generate new structural models. Since the quality of the final structure is influenced by the initial structure, the optimization process is carried out 20 times, each with a random structure as the start point. The optimized structure with the lowest energy is selected as the final predicted structure of a dimer.

### Markov Chain Monte Carlo Optimization

We apply a Rosetta protocol in pyRosetta based on Metropolis-Hasting sampling^49^ to implement a Markov Chain Monte Carlo (MC) optimization to reconstruct complex structures according to the Boltzmann distribution. An initial conformation of a dimer is generated in the same way as in the GD algorithm. Starting from the initial conformation, a low-resolution rigid-body search is employed to rotate and translate one chain around the surface of the other chain to generate new structures in the MC optimization. 500 Monte Carlo moves are attempted. Each move is accepted or rejected based on the standard Metropolis acceptance criterion^50^.

After the low-resolution search, back-bone and side-chain conformations are further optimized with the Newton minimization method in a high-resolution refinement process, in which the gradient of the scoring function dictates the direction of the starting point in the rigid-body translation/rotation space. This minimization process is repeated 50 times to detect the local minimum of the energy function that may have similar performance as the global minimum^6^.

We implement the MC method above using high-resolution and low-resolution docking protocols in RosettaDock to optimize the same energy function used in the GD method. Low-resolution docking is performed using the DockingLowRes protocol, whereas DockMCMProtocol is used to perform high-resolution docking. For a dimer, *10*^*5*^ to *10*^*7*^ rounds of MC optimization with different initial conformations are executed to generate structural models. At the end,*10*^*5*^ to *10*^*7*^ models are generated, among which the model with the lowest energy is selected as the final prediction.

### Simulated Annealing Optimization Based on Crystallography and NMR System (CNS)

This structure optimization method, Con_Complex^41^ in the DeepComplex package, is implemented on top of the Crystallography and NMR System (CNS)^51,52^ that uses a simulated annealing protocol to search for quaternary structures that satisfy inter-chain contacts^46^. This method takes the PDB files of monomers (protein chains) in a protein multimer (e.g., homodimer) and the true or predicted inter-protein contacts as input to reconstruct the structure of the multimer without altering the shape of the structure of the monomer. The inter-protein contacts are converted into distance restraints used by CNS. This process generates 100 structural models and then picks 5 models with lowest CNS energy. It is worth noting that this method can handle the reconstruction of the quaternary structure of any multimer consisting of multiple identical or different chains. Because inter-chain contacts are the main restraints to guide structure modeling, the performance of this method mostly depends on the quality of the inter-protein contact predictions.

## Supporting information

Supplemental Data for Distance-based Reconstruction of Protein Quaternary Structures from Inter-Chain Contacts

## Data Availability

The source code of the methods and test data sets are available at: https://github.com/jianlin-cheng/DeepComplex2

## Acknowledgements

Research reported in this publication was supported in part by Department of Energy grants (DEAR0001213, DE-SC0020400 and DESC0021303), two NSF grants (DBI 1759934 and IIS1763246), and an NIH grant (R01GM093123).

## Author Contributions

JC conceived the project. ES and JC designed the method. ES, FQ, and RR implemented the method and collected the results. ES, FQ, RR, and JC analyzed the results. ES, FQ, RR, and JC wrote and manuscript.

