## Supplemental Data for Distance-based Reconstruction of Protein Quaternary Structures from Inter-Chain Contacts for "Distance-based Reconstruction of Protein Quaternary Structures from Inter-Chain Contacts"

**Supplementary Figures**

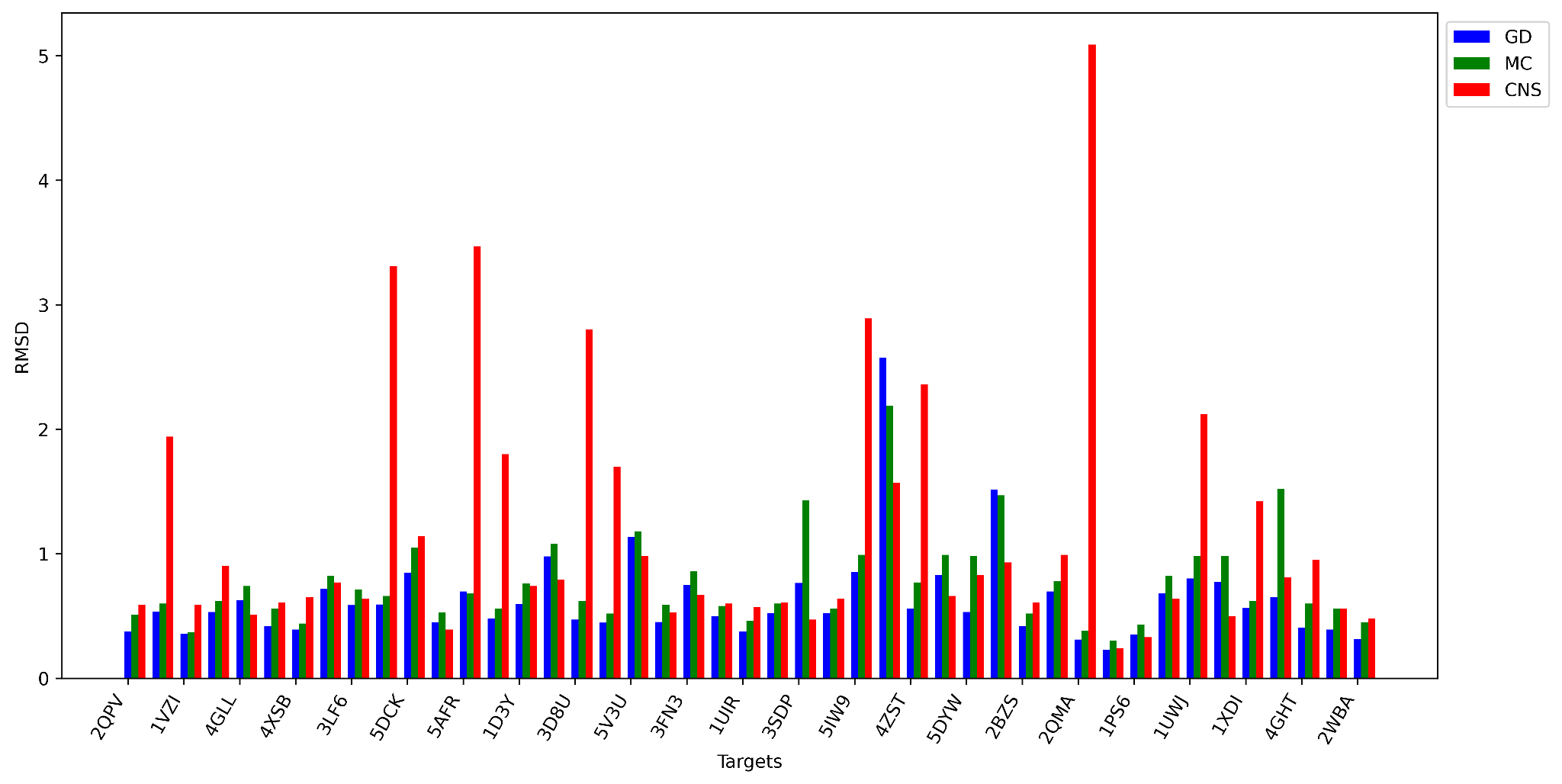

**Figure S1**. RMSD of GD, MC, and CNS on a dataset of 44 homodimers with known inter-protein contacts. The average RMSD of GD, MC, and CNS is 0.63, 0.76, and 1.16.

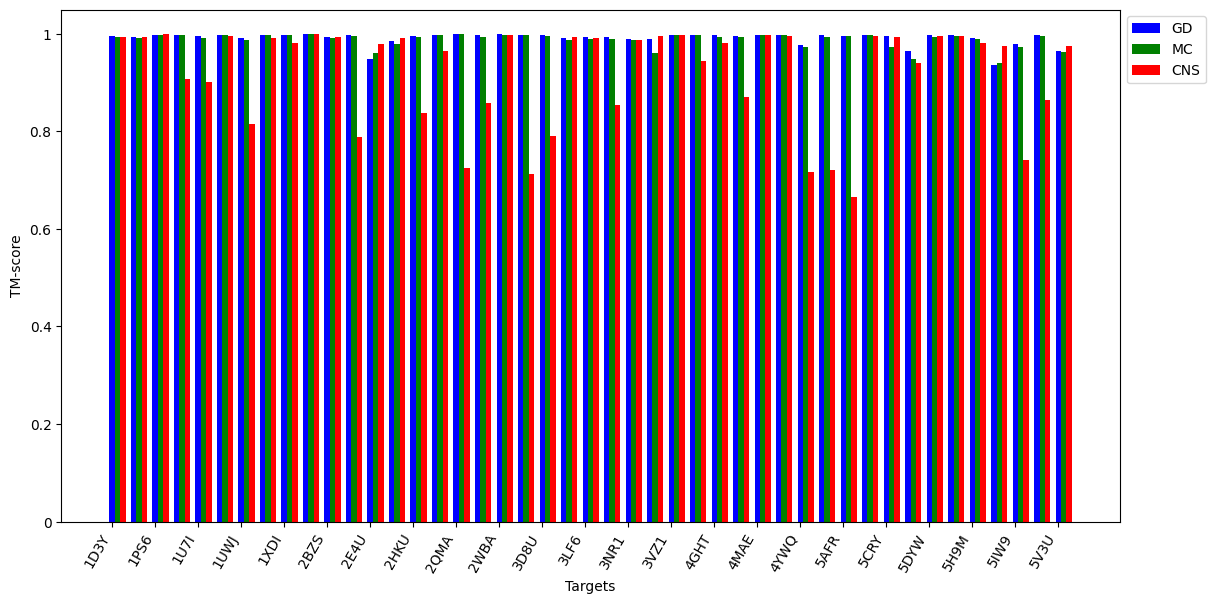

**Figure S2**. TM-score of GD, MC, and CNS on a dataset of 44 protein complexes with known inter-protein contacts. The average TM-score of GD, MC, and CNS is 0.99, 0.98, and 0.91.

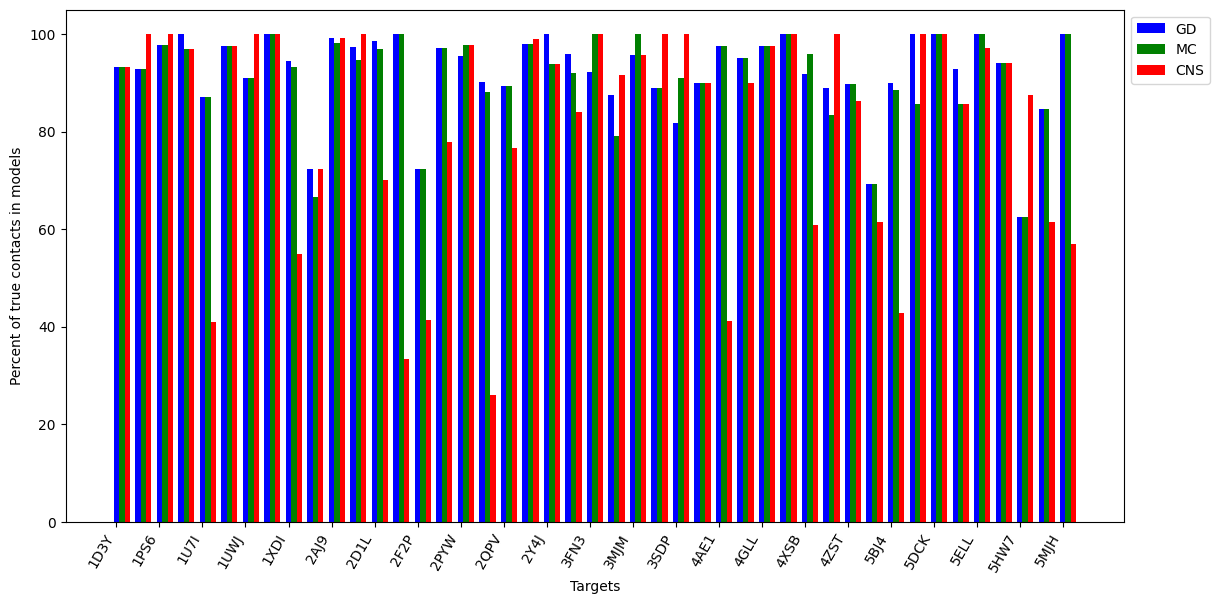

**Figure S3**. The f_nat of GD, MC, and CNS on a dataset of 44 homodimers with known inter-protein contacts. The average f_nat of GD, MC, and CNS is 92.19, 91.39, and 82.49.

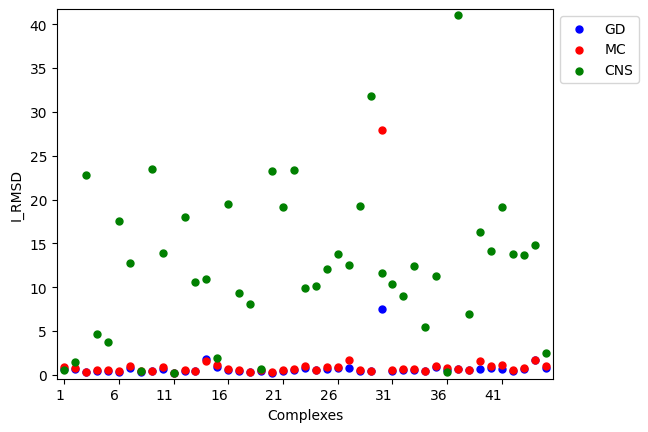

**Figure S4**. I_RMSD of GD, MC, and CNS on a dataset of 44 homodimers with known inter-protein contacts. The average I_RMSD of GD, MC, and CNS is 0.77, 1.35, and 12.46.

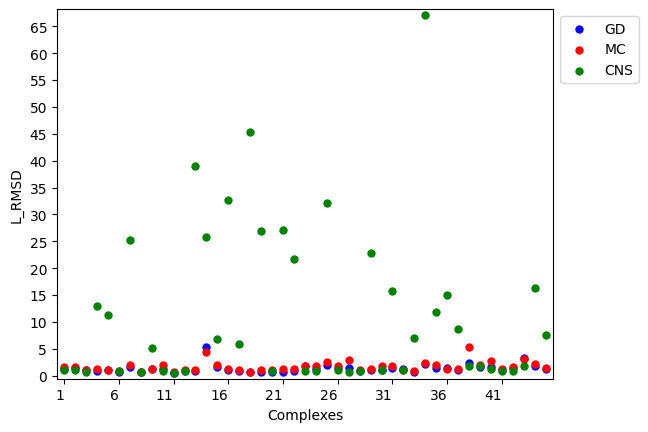

**Figure S5**. L_RMSD of GD, MC, and CNS on a dataset of 44 homodimers with known inter-protein contacts. The average L_RMSD of GD, MC, and CNS is 1.38, 1.7, and 11.18.

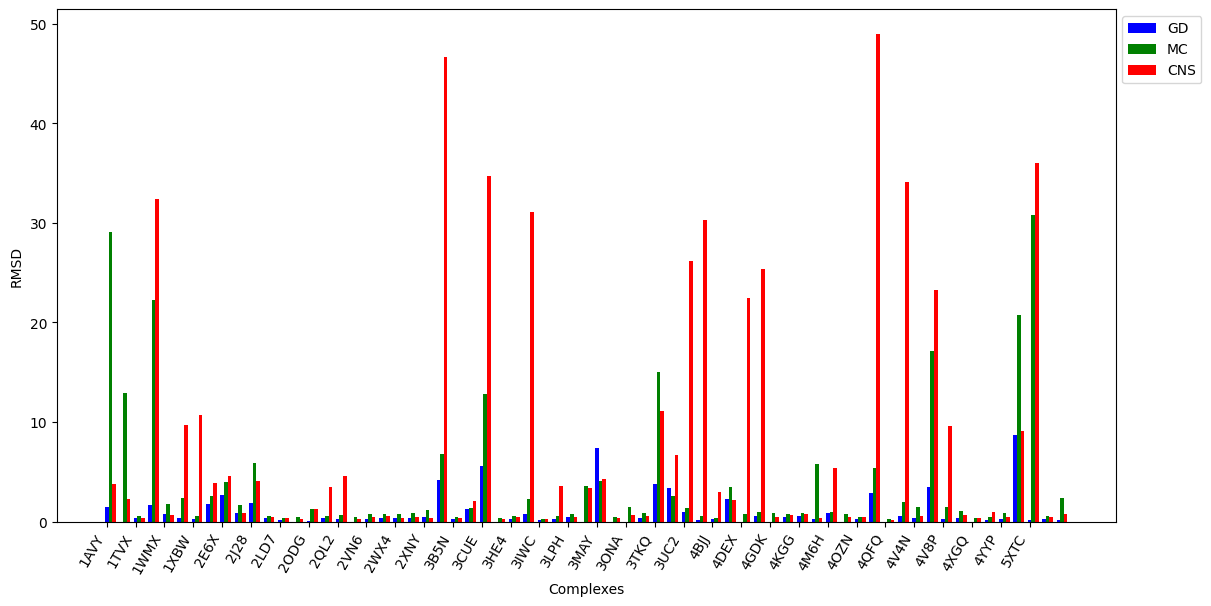

**Figure S6**. RMSD of GD, MC, and CNS on 73 heterodimers with known inter-protein contacts. The average RMSD of GD, MC, and CNS is 1.23, 4.76, and 7.7.

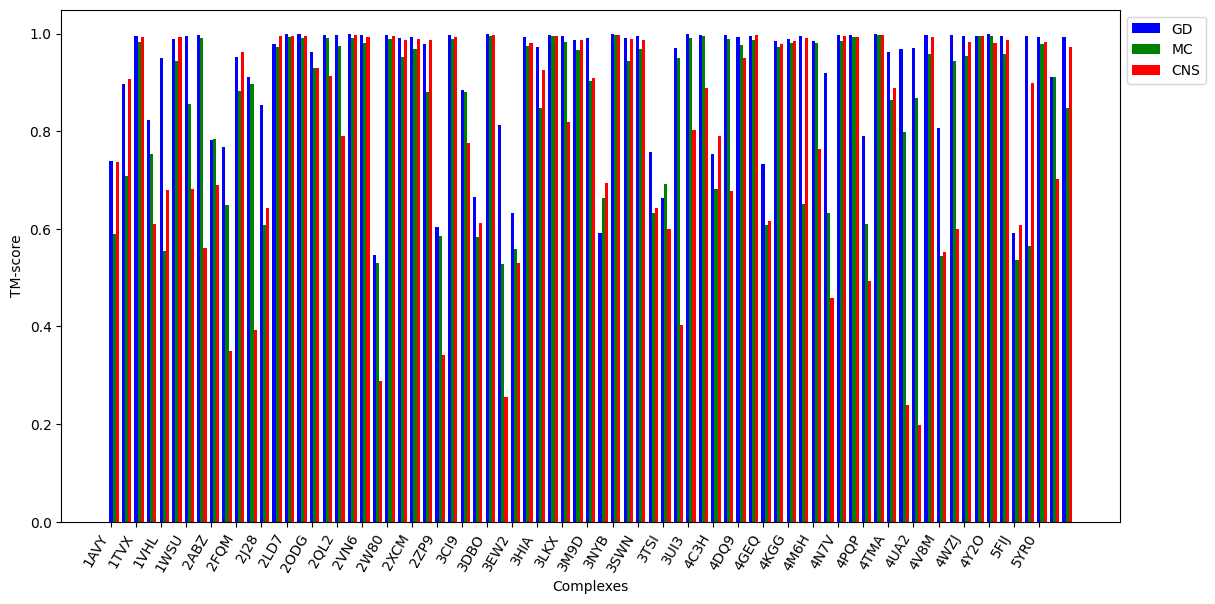

**Figure S7**. TM-score of GD, MC, and CNS on 73 heterodimers with known inter-protein contacts. The average TM-score of GD, MC, and CNS is 0.92, 0.85, and 0.79.

**Supplementary Tables**

**Table S1**. TM-score, RMSD, f_nat, I_RMSD, and L_RMSD of the models that GD reconstructed for each of 44 homodimers from true inter-chain contacts.

| **Target** | **Number of true contacts** | **Length of chain A** | **Length of chain B** | **TM-score** | **RMSD** | **f_nat (%)** | **I_RMSD** | **L_RMSD** |
| --- | --- | --- | --- | --- | --- | --- | --- | --- |
| **2QMA** | 621 | 444 | 440 | 0.999 | 0.31 | 90.2 | 0.325 | 0.619 |
| **2E4U** | 52 | 512 | 514 | 0.948 | 2.576 | 100 | 1.85 | 5.314 |
| **5DCK** | 53 | 71 | 72 | 0.965 | 0.846 | 100 | 0.754 | 1.674 |
| **5CRY** | 59 | 348 | 348 | 0.995 | 0.652 | 100 | 0.682 | 2.315 |
| **5IW9** | 67 | 123 | 122 | 0.979 | 0.853 | 84.6 | 0.842 | 1.727 |
| **4GHT** | 77 | 181 | 181 | 0.997 | 0.407 | 95 | 0.41 | 1.128 |
| **3SDP** | 78 | 186 | 186 | 0.988 | 0.764 | 81.8 | 0.813 | 1.438 |
| **5V3U** | 83 | 131 | 123 | 0.965 | 1.135 | 90.9 | 1.12 | 2.64 |
| **5DYW** | 84 | 527 | 525 | 0.998 | 0.532 | 92.9 | 0.62 | 1.682 |
| **5AFR** | 88 | 327 | 325 | 0.994 | 0.696 | 69.19 | 0.681 | 1.349 |
| **4YWQ** | 89 | 147 | 146 | 0.977 | 0.979 | 88.9 | 0.847 | 2.19 |
| **3FN3** | 94 | 215 | 211 | 0.99 | 0.748 | 92.3 | 0.84 | 1.737 |
| **2Y4J** | 99 | 377 | 377 | 0.998 | 0.391 | 100 | 0.398 | 0.788 |
| **5H9M** | 103 | 190 | 189 | 0.99 | 0.696 | 94.1 | 0.642 | 1.312 |
| **3D8U** | 107 | 260 | 266 | 0.997 | 0.471 | 96 | 0.719 | 1.39 |
| **3LF6** | 110 | 154 | 157 | 0.992 | 0.588 | 87.5 | 0.526 | 0.949 |
| **1UWJ** | 111 | 264 | 263 | 0.99 | 0.802 | 90.9 | 0.564 | 1.247 |
| **1JCZ** | 116 | 260 | 260 | 0.993 | 0.683 | 92.9 | 0.833 | 1.603 |
| **3NR1** | 130 | 178 | 178 | 0.989 | 0.717 | 88.9 | 0.674 | 1.43 |
| **3MJM** | 132 | 343 | 342 | 0.993 | 0.773 | 95.7 | 0.748 | 1.512 |
| **4GLL** | 139 | 307 | 306 | 0.995 | 0.625 | 97.5 | 0.657 | 2.08 |
| **5ELL** | 142 | 231 | 235 | 0.996 | 0.499 | 100 | 0.558 | 1.423 |
| **1PS6** | 144 | 328 | 328 | 0.998 | 0.348 | 97.7 | 0.438 | 1.108 |
| **4ZST** | 146 | 328 | 328 | 0.996 | 0.56 | 89.7 | 0.281 | 0.999 |
| **2HKU** | 152 | 188 | 182 | 0.994 | 0.534 | 97.2 | 0.569 | 1.358 |
| **2F2P** | 155 | 169 | 169 | 0.985 | 0.827 | 72.39 | 0.603 | 1.047 |
| **4MAE** | 161 | 577 | 577 | 0.998 | 0.522 | 100 | 0.924 | 1.598 |
| **1U7I** | 164 | 130 | 129 | 0.994 | 0.48 | 87.2 | 0.601 | 1.219 |
| **1SOX** | 167 | 463 | 458 | 0.998 | 0.452 | 100 | 0.492 | 0.975 |
| **2QPV** | 169 | 128 | 128 | 0.996 | 0.375 | 89.4 | 0.46 | 0.964 |
| **1VZI** | 174 | 125 | 125 | 0.996 | 0.355 | 100 | 0.417 | 0.729 |
| **1UIR** | 179 | 309 | 313 | 0.998 | 0.376 | 97.6 | 0.373 | 0.717 |
| **2PYW** | 197 | 417 | 411 | 0.998 | 0.448 | 95.6 | 0.377 | 0.763 |
| **2BZS** | 207 | 230 | 228 | 0.993 | 0.419 | 97.3 | 0.45 | 0.949 |
| **2D1L** | 236 | 240 | 222 | 0.997 | 0.447 | 98.5 | 0.441 | 0.859 |
| **1D3Y** | 250 | 289 | 290 | 0.995 | 0.597 | 93.3 | 0.423 | 0.914 |
| **3VZ1** | 273 | 452 | 452 | 0.998 | 0.417 | 90.0 | 0.671 | 1.341 |
| **5MJH** | 320 | 368 | 368 | 0.996 | 0.593 | 100 | 0.418 | 0.844 |
| **4XSB** | 323 | 340 | 343 | 0.998 | 0.39 | 91.9 | 0.65 | 1.261 |
| **5BJ4** | 340 | 366 | 366 | 0.997 | 0.524 | 90 | 0.403 | 0.784 |
| **2WBA** | 347 | 489 | 489 | 0.999 | 0.313 | 98 | 0.535 | 1.049 |
| **1XDI** | 352 | 459 | 459 | 0.997 | 0.562 | 94.5 | 0.249 | 0.747 |
| **2AJ9** | 364 | 334 | 334 | 0.999 | 0.228 | 99.1 | 0.48 | 1.225 |
| **4AE1** | 398 | 501 | 501 | 0.997 | 0.531 | 97.5 | 0.204 | 0.446 |
| **5HW7** | 39 | 122 | 119 | 0.936 | 1.515 | 62.5 | 0.419 | 1.061 |

**Table S2**. TM-score, RMSD, f_nat, I_RMSD, and L_RMSD of structural models reconstructed by GD for 73 heterodimers in the Hetero73 dataset using true/native inter-chain contacts as input.

| **Target** | **Chains** | **Length of chain 1** | **Length of chain 2** | **Number of true contacts** | **TM-score** | **RMSD** | **f_nat** | **I_RMSD** | **L_RMSD** |
| --- | --- | --- | --- | --- | --- | --- | --- | --- | --- |
| **1AVY** | A, B | 68 | 54 | 19 | 0.74 | 1.47 | 92.9 | 0 | 6.91 |
| **1IS7** | A, K | 194 | 84 | 10 | 0.9 | 0 | 85.7 | 1.61 | 4.97 |
| **1TVX** | A, B | 64 | 71 | 86 | 0.99 | 0.33 | 100 | 0.35 | 0.66 |
| **1VCH** | B, E | 170 | 152 | 19 | 0.82 | 3.33 | 42.9 | 1.76 | 9.22 |
| **1VHL** | B, C | 208 | 183 | 22 | 0.95 | 1.65 | 72.7 | 1.53 | 4.04 |
| **1WMX** | A, B | 173 | 195 | 61 | 0.99 | 0.74 | 96.89 | 0.49 | 1.75 |
| **1WSU** | C, D | 102 | 121 | 49 | 0.99 | 0.39 | 100 | 0.38 | 1.21 |
| **1XBW** | A, B | 99 | 96 | 149 | 1 | 0.26 | 97.7 | 0.25 | 0.63 |
| **2ABZ** | C, D | 62 | 46 | 13 | 0.78 | 1.74 | 54.5 | 2 | 5.06 |
| **2E6X** | C, D | 56 | 66 | 23 | 0.77 | 2.64 | 85.7 | 0 | 5.38 |
| **2FQM** | A, D | 65 | 72 | 21 | 0.95 | 0.84 | 100 | 0.76 | 3.57 |
| **2IS5** | A, D | 156 | 143 | 43 | 0.91 | 2.01 | 61.9 | 1.88 | 4.04 |
| **2J28** | 1, 3 | 54 | 64 | 11 | 0.85 | 1.82 | 83.3 | 0.93 | 6.05 |
| **2JG8** | A, B | 132 | 129 | 119 | 0.98 | 0.32 | 98.6 | 0.32 | 0.67 |
| **2LD7** | A, B | 94 | 75 | 153 | 1 | 0.18 | 95.7 | 0.18 | 0.34 |
| **2MJF** | A, B | 40 | 95 | 132 | 1 | 0 | 97.8 | 0.16 | 0.3 |
| **2ODG** | A, C | 89 | 47 | 30 | 0.96 | 0.07 | 78.6 | 0 | 1.65 |
| **2P7M** | A, B | 127 | 122 | 199 | 1 | 0.33 | 97 | 0.33 | 0.65 |
| **2QL2** | A, B | 56 | 59 | 79 | 1 | 0.25 | 94.69 | 0.23 | 0.58 |
| **2ROZ** | A, B | 32 | 136 | 101 | 1 | 0 | 100 | 0.21 | 0.46 |
| **2VN6** | A, B | 151 | 64 | 80 | 1 | 0.27 | 100 | 0.3 | 0.61 |
| **2W80** | A, D | 123 | 244 | 145 | 1 | 0.39 | 100 | 0.42 | 0.87 |
| **2WX4** | B, C | 41 | 43 | 61 | 0.99 | 0.3 | 100 | 0.3 | 0.6 |
| **2XCM** | C, E | 92 | 74 | 57 | 0.99 | 0.4 | 100 | 0.37 | 0.81 |
| **2XNY** | M, N | 37 | 36 | 77 | 0.98 | 0.44 | 83.6 | 0.43 | 0.87 |
| **2ZP9** | E, I | 49 | 39 | 4 | 0.6 | 4.17 | 100 | 3.06 | 16.47 |
| **3B5N** | B, C | 69 | 70 | 149 | 1 | 0.23 | 100 | 0.23 | 0.45 |
| **3CI9** | A, B | 44 | 45 | 20 | 0.88 | 1.28 | 100 | 0.54 | 4.62 |
| **3CUE** | B, Q | 167 | 188 | 5 | 0.67 | 5.6 | 50 | 0.98 | 23.58 |
| **3DBO** | A, B | 34 | 126 | 171 | 1 | 0 | 98.1 | 0.1 | 0.44 |
| **3ERM** | D, E | 63 | 56 | 16 | 0.81 | 2.14 | 81.8 | 0.43 | 6.61 |
| **3EW2** | C, F | 124 | 119 | 10 | 0.63 | 5.36 | 85.7 | 2.06 | 20.79 |
| **3HE4** | A, B | 45 | 44 | 90 | 0.99 | 0.28 | 100 | 0.25 | 0.53 |
| **3HIA** | B, C | 83 | 74 | 38 | 0.97 | 0.8 | 88.5 | 0.69 | 1.72 |
| **3IWC** | A, B | 58 | 61 | 255 | 1 | 0.18 | 96.89 | 0.18 | 0.36 |
| **3LKX** | A, B | 65 | 53 | 123 | 1 | 0.28 | 100 | 0.28 | 0.54 |
| **3LPH** | A, B | 62 | 55 | 65 | 0.99 | 0.44 | 100 | 0.44 | 0.97 |
| **3M9D** | A, G | 186 | 31 | 29 | 0.99 | 0 | 90.9 | 0.71 | 1.55 |
| **3MAY** | A, C | 86 | 97 | 6 | 0.59 | 7.39 | 83.3 | 2.78 | 15.12 |
| **3NYB** | A, B | 323 | 64 | 173 | 1 | 0 | 99.2 | 0.27 | 0.57 |
| **3ONA** | A, B | 158 | 66 | 61 | 0.99 | 0 | 97.1 | 0.46 | 1.22 |
| **3SWN** | R, S | 76 | 72 | 98 | 1 | 0.32 | 100 | 0.32 | 0.66 |
| **3TKQ** | A, E | 191 | 166 | 5 | 0.76 | 3.8 | 50 | 4.74 | 10.04 |
| **3UC2** | B, D | 125 | 109 | 43 | 0.97 | 0.98 | 88.5 | 0.76 | 2.63 |
| **3UI3** | A, B | 142 | 102 | 177 | 1 | 0.17 | 99.1 | 0.17 | 0.39 |
| **4BJJ** | A, B | 106 | 85 | 192 | 1 | 0.28 | 96.5 | 0.29 | 0.56 |
| **4C3H** | J, L | 69 | 45 | 18 | 0.75 | 2.26 | 90 | 0 | 7.34 |
| **4DEX** | A, B | 289 | 45 | 93 | 1 | 0 | 96.2 | 0.38 | 1.03 |
| **4DQ9** | A, B | 149 | 141 | 66 | 0.99 | 0.51 | 98 | 0.49 | 1.02 |
| **4GDK** | A, B | 88 | 267 | 71 | 0.99 | 0 | 95.8 | 0.37 | 1.96 |
| **4GEQ** | B, C | 58 | 90 | 6 | 0.73 | 2.61 | 50 | 1.57 | 15.37 |
| **4K12** | A, B | 64 | 82 | 57 | 0.98 | 0.5 | 100 | 0.45 | 1.71 |
| **4KGG** | D, A | 163 | 141 | 66 | 0.99 | 0.57 | 96.2 | 0.56 | 1.16 |
| **4M3L** | A, D | 60 | 53 | 59 | 0.99 | 0.28 | 100 | 0.3 | 0.67 |
| **4M6H** | A, B | 190 | 162 | 66 | 0.99 | 0.84 | 97.1 | 0.59 | 2.14 |
| **4M77** | H, J | 85 | 72 | 23 | 0.92 | 1.46 | 100 | 0.56 | 3.81 |
| **4N7V** | A, C | 222 | 33 | 98 | 1 | 0 | 100 | 0.2 | 0.76 |
| **4OZN** | A, B | 116 | 104 | 124 | 1 | 0.28 | 98.8 | 0.29 | 0.68 |
| **4PQP** | A, D | 102 | 97 | 17 | 0.79 | 2.89 | 64.3 | 1.24 | 8.39 |
| **4QFQ** | A, B | 101 | 35 | 250 | 1 | 0 | 98.4 | 0 | 0.29 |
| **4TMA** | I, J | 47 | 57 | 42 | 0.96 | 0.6 | 93.5 | 0.57 | 1.83 |
| **4U3Q** | A, B | 93 | 99 | 17 | 0.97 | 0.94 | 81.8 | 0.94 | 2.62 |
| **4UA2** | A, H | 115 | 103 | 27 | 0.97 | 0.95 | 94.1 | 0.59 | 2.08 |
| **4V4N** | T, W | 215 | 135 | 107 | 1 | 0.31 | 100 | 0.35 | 0.85 |
| **4V8P** | K, M | 108 | 143 | 65 | 1 | 0.28 | 100 | 0.28 | 0.95 |
| **4WZJ** | L, M | 79 | 79 | 98 | 0.99 | 0.34 | 98.1 | 0.32 | 0.64 |
| **4XGQ** | A, B | 132 | 30 | 90 | 1 | 0 | 98.5 | 0.36 | 0.77 |
| **4Y2O** | A, B | 211 | 142 | 170 | 1 | 0.19 | 98 | 0.2 | 0.44 |
| **4YYP** | A, B | 87 | 32 | 86 | 0.99 | 0.3 | 100 | 0.33 | 0.71 |
| **5FIJ** | S, T | 167 | 174 | 7 | 0.59 | 8.68 | 11.4 | 0 | 36.13 |
| **5XTC** | B, V | 124 | 111 | 64 | 0.99 | 0.17 | 97.1 | 0.44 | 0.61 |
| **5YR0** | A, B | 48 | 44 | 75 | 0.99 | 0.3 | 100 | 0.29 | 0.58 |
| **6UMM** | D, I | 81 | 61 | 23 | 0.99 | 0.11 | 100 | 0.48 | 0.68 |

Table S3. Average TM-score, RMSD, f_nat, I_RMSD, and L_RMSD) of GD on 32 heterodimers in the Std32 dataset using true contacts as input.

| **Target** | **Length of chain 1** | **Length of chain 2** | **Chains** | **Number of true contacts** | **TM-score** | **RMSD** | **f_nat** | **I_RMSD** | **L_RMSD** |
| --- | --- | --- | --- | --- | --- | --- | --- | --- | --- |
| **1W85** | 358 | 324 | A, B | 185 | 1 | 0.11 | 100 | 0.1 | 0.22 |
| **1EFP** | 307 | 246 | A, B | 317 | 1 | 0.07 | 97.8 | 0.08 | 0.18 |
| **1I1Q** | 512 | 186 | A, B | 153 | 1 | 0.08 | 100 | 0.08 | 0.18 |
| **2Y69** | 227 | 259 | B, C | 2 | 0.54 | 13.88 | 50 | 10 | 11.01 |
| **3MML** | 285 | 207 | A, B | 146 | 1 | 0.12 | 96.3 | 0.12 | 0.38 |
| **2VPZ** | 734 | 193 | A, B | 188 | 1 | 0.19 | 94.1 | 0.24 | 0.58 |
| **1TYG** | 65 | 242 | B, A | 114 | 1 | 0.05 | 98.7 | 0.08 | 0.26 |
| **3RPF** | 143 | 72 | A, C | 5 | 0.82 | 3.49 | 100 | 1.15 | 7.44 |
| **1EP3** | 311 | 261 | A, B | 133 | 1 | 0.05 | 100 | 0.04 | 0.12 |
| **2NU9** | 285 | 385 | A, B | 190 | 1 | 0.11 | 98.6 | 0.11 | 0.21 |
| **3RRL** | 227 | 197 | A, B | 194 | 1 | 0.14 | 99.3 | 0.14 | 0.45 |
| **3IP4** | 485 | 482 | A, B | 183 | 1 | 0.12 | 98.3 | 0.02 | 0.24 |
| **1RM6** | 761 | 323 | A, B | 129 | 1 | 0.18 | 100 | 0.1 | 0.28 |
| **2D1P** | 119 | 95 | B, C | 60 | 1 | 0.12 | 97.3 | 0.1 | 0.33 |
| **4HR7** | 443 | 80 | A, B | 36 | 0.99 | 0.78 | 100 | 0.35 | 2.2 |
| **2ONK** | 240 | 252 | A, C | 92 | 1 | 0.32 | 92.6 | 0 | 0.9 |
| **3A0R** | 334 | 113 | A, B | 60 | 1 | 0.28 | 100 | 0.06 | 0.71 |
| **1B70** | 265 | 775 | A, B | 414 | 1 | 0.07 | 95.7 | 0.08 | 0.23 |
| **1QOP** | 265 | 390 | A, B | 157 | 1 | 0.15 | 99 | 0.15 | 0.32 |
| **2WDQ** | 121 | 105 | C, D | 77 | 1 | 0.12 | 100 | 0.12 | 0.27 |
| **1BXR** | 1073 | 379 | A, B | 258 | 1 | 0.07 | 98.7 | 0.07 | 0.17 |
| **3G5O** | 92 | 81 | A, B | 143 | 1 | 0.05 | 97.8 | 0.05 | 0.08 |
| **3OAA** | 138 | 284 | H, G | 240 | 1 | 0.11 | 98.3 | 0.11 | 0.33 |
| **3PNL** | 356 | 211 | A, B | 148 | 1 | 0.2 | 100 | 0.19 | 0.55 |
| **1ZUN** | 196 | 382 | A, B | 251 | 1 | 0.14 | 100 | 0.15 | 0.45 |
| **1IXR** | 135 | 308 | A, C | 0 | 0.69 | 29.9 | 0 | 29 | 92.99 |
| **1W85** | 358 | 324 | A, B | 185 | 1 | 0.11 | 100 | 0.1 | 0.22 |
| **1EFP** | 307 | 246 | A, B | 317 | 1 | 0.07 | 97.8 | 0.08 | 0.18 |
| **1I1Q** | 512 | 186 | A, B | 153 | 1 | 0.08 | 100 | 0.08 | 0.18 |
| **2Y69** | 227 | 259 | B, C | 2 | 0.54 | 13.88 | 50 | 10 | 11.01 |
| **3MML** | 285 | 207 | A, B | 146 | 1 | 0.12 | 96.3 | 0.12 | 0.38 |
| **2VPZ** | 734 | 193 | A, B | 188 | 1 | 0.19 | 94.1 | 0.24 | 0.58 |

Table S4. Detailed results of GD on Set A with predicted inter-chain contacts as input.

| **Target name** | **Length of chain A** | **Length of chain B** | **Number of predicted interchain contacts** | **Precision of predicted interchain contacts (%)** | **Recall of predicted inter-chain contacts (%)** | **TM-score** | **RMSD** | **f_nat** | **I_RMSD** | **L_RMSD** |
| --- | --- | --- | --- | --- | --- | --- | --- | --- | --- | --- |
| **2XBQ** | 105 | 105 | 26 | 0.0 | 0.0 | 0.5 | 14.74 | 0.0 | 14.544 | 43.547 |
| **1Z9Z** | 60 | 60 | 32 | 0.0 | 0.0 | 0.55 | 11.27 | 0.0 | 13.191 | 20.364 |
| **1A19** | 89 | 89 | 26 | 0.0 | 0.0 | 0.5 | 14.96 | 0.0 | 14.706 | 39.503 |
| **1YH8** | 266 | 266 | 54 | 0.0 | 0.0 | 0.5 | 18.47 | 0.0 | 17.312 | 58.735 |
| **3N8E** | 159 | 159 | 53 | 0.0 | 0.0 | 0.51 | 21.39 | 0.0 | 22.79 | 44.298 |
| **5LLJ** | 57 | 57 | 41 | 3.12 | 8.0 | 0.53 | 13.20 | 0.0 | 9.685 | 22.742 |
| **2FU4** | 81 | 81 | 29 | 0.0 | 0.0 | 0.5 | 17.39 | 0.0 | 11.888 | 38.914 |
| **2PL7** | 67 | 67 | 25 | 1.85 | 3.33 | 0.51 | 10.19 | 0.0 | 9.634 | 28.831 |
| **4E83** | 31 | 31 | 41 | 5.08 | 9.68 | 0.55 | 9.36 | 12.5 | 6.373 | 13.754 |
| **5UZX** | 231 | 231 | 48 | 0.0 | 0.0 | 0.5 | 22.67 | 0.0 | 24.152 | 66.521 |
| **1A2D** | 130 | 130 | 53 | 2.35 | 5.88 | 0.67 | 6.44 | 10 | 6.57 | 11.312 |
| **1RRG** | 177 | 177 | 62 | 0.0 | 0.0 | 0.64 | 8.71 | 0.0 | 7.635 | 16.947 |
| **3JSL** | 308 | 308 | 29 | 0.0 | 0.0 | 0.5 | 23.37 | 0.0 | 24.824 | 55.067 |
| **3LO2** | 30 | 30 | 43 | 37.5 | 50 | 0.86 | 1.1 | 75 | 1.048 | 2.201 |
| **4Q1R** | 130 | 130 | 24 | 46.34 | 52.78 | 0.98 | 0.62 | 84.6 | 0.614 | 1.78 |
| **1VH9** | 138 | 138 | 82 | 3.48 | 10.81 | 0.5 | 13.97 | 16.7 | 13.865 | 35.895 |
| **1D8U** | 165 | 165 | 48 | 8.86 | 18.42 | 0.6 | 7.48 | 0.0 | 7.45 | 17.776 |
| **4GA9** | 134 | 134 | 58 | 54.69 | 85.37 | 0.9 | 2.06 | 57.1 | 2.083 | 3.787 |
| **1IU8** | 206 | 206 | 38 | 8.97 | 14.89 | 0.61 | 18.85 | 0.0 | 16.81 | 26.005 |
| **2R74** | 142 | 142 | 61 | 0.0 | 0.0 | 0.5 | 15.99 | 0.0 | 15.796 | 45.492 |
| **5C39** | 51 | 51 | 90 | 34.67 | 52 | 0.99 | 0.26 | 92.3 | 0.196 | 0.518 |
| **1F86** | 115 | 115 | 82 | 32.67 | 63.46 | 0.99 | 0.36 | 100 | 0.397 | 0.724 |
| **2ZWM** | 120 | 120 | 21 | 1.39 | 1.92 | 0.62 | 11.58 | 0.0 | 11.444 | 18.365 |
| **1M0U** | 203 | 203 | 94 | 46.08 | 85.45 | 0.99 | 0.7 | 100 | 0.766 | 1.427 |
| **1PD3** | 54 | 54 | 28 | 1.22 | 1.82 | 0.94 | 0.94 | 20.8 | 1.2 | 1.98 |
| **2D4G** | 165 | 165 | 98 | 2.68 | 7.27 | 0.6 | 19.53 | 0.0 | 12.094 | 28.873 |
| **3F08** | 135 | 135 | 113 | 0.58 | 1.67 | 0.53 | 16.71 | 0.0 | 17.982 | 32.181 |
| **2CC3** | 144 | 144 | 28 | 2.3 | 3.28 | 0.62 | 9.01 | 0.0 | 7.978 | 15.46 |
| **1GNW** | 210 | 210 | 34 | 28 | 33.87 | 0.99 | 0.54 | 60.9 | 0.562 | 1.046 |
| **2CCY** | 127 | 127 | 53 | 11.65 | 19.35 | 0.64 | 17.74 | 0.0 | 19.329 | 25.58 |
| **5F5X** | 333 | 333 | 30 | 13.41 | 17.46 | 0.7 | 5.38 | 0.0 | 5.183 | 11.139 |
| **5JYB** | 344 | 344 | 37 | 14.77 | 20.31 | 0.97 | 1.61 | 11.8 | 1.668 | 4.962 |
| **1EOG** | 208 | 208 | 99 | 51.85 | 86.15 | 0.99 | 0.29 | 96.2 | 0.311 | 0.629 |
| **1MK4** | 157 | 157 | 24 | 0.0 | 0.0 | 0.6 | 12.29 | 0.0 | 12.115 | 20.927 |
| **1HNB** | 217 | 217 | 58 | 40.45 | 53.73 | 0.99 | 0.61 | 81 | 0.526 | 1.348 |
| **2CVI** | 83 | 83 | 40 | 5.94 | 8.96 | 0.62 | 6.19 | 0.0 | 6.193 | 11.909 |
| **1V8F** | 276 | 276 | 35 | 17.05 | 22.06 | 0.5 | 26.92 | 0.0 | 21.932 | 56.044 |
| **1YQ1** | 198 | 198 | 51 | 36.78 | 47.06 | 1 | 0.11 | 95.7 | 0.118 | 0.214 |
| **3BBH** | 204 | 204 | 22 | 8.43 | 10.29 | 0.92 | 2.11 | 29.4 | 2.423 | 4.107 |
| **3RHU** | 141 | 141 | 49 | 0.0 | 0.0 | 0.5 | 21.20 | 0.0 | 22.02 | 57.47 |

Table S5. Detailed results of GD on Set B with predicted inter-chain contacts as input.

| **Target name** | **Length of chain A** | **Length of chain B** | **Number of predicted interchain contacts** | **Precision of predicted interchain contacts (%)** | **Recall of predicted inter-chain contacts (%)** | **TM-score** | **RMSD** | **f_nat** | **I_RMSD** | **L_RMSD** |
| --- | --- | --- | --- | --- | --- | --- | --- | --- | --- | --- |
| **1LBK** | 208 | 208 | 94 | 52.34 | 81.16 | 0.99 | 0.26 | 96.6 | 0.27 | 0.56 |
| **2YYB** | 242 | 242 | 31 | 3.06 | 4.29 | 0.5 | 25.28 | 0.0 | 23.48 | 53.45 |
| **1T92** | 108 | 108 | 29 | 3.06 | 4.17 | 0.64 | 5.97 | 0.0 | 5.58 | 14.56 |
| **2QY6** | 244 | 244 | 28 | 0.0 | 0.0 | 0.5 | 25.84 | 0.0 | 21.64 | 72.11 |
| **1ML6** | 219 | 219 | 72 | 51.04 | 67.12 | 1 | 0.1 | 96.6 | 0.12 | 0.22 |
| **5AIF** | 124 | 124 | 22 | 6.52 | 7.89 | 0.94 | 1.36 | 24 | 1.38 | 2.80 |
| **2YR1** | 257 | 257 | 26 | 8.42 | 10.39 | 0.96 | 1.51 | 22.2 | 1.59 | 4.12 |
| **1B48** | 221 | 221 | 68 | 55.32 | 66.67 | 0.99 | 0.24 | 100 | 0.26 | 0.5 |
| **3F1V** | 366 | 366 | 39 | 27.17 | 32.05 | 0.99 | 0.77 | 56 | 0.76 | 2.47 |
| **3KXO** | 198 | 198 | 86 | 47.75 | 67.95 | 0.99 | 0.28 | 100 | 0.29 | 0.53 |
| **1ECS** | 120 | 120 | 94 | 25.38 | 44.3 | 0.97 | 0.91 | 71.4 | 0.80 | 1.90 |
| **3EE2** | 198 | 198 | 90 | 46.69 | 68.35 | 0.99 | 0.33 | 92 | 0.37 | 0.74 |
| **3WVA** | 163 | 163 | 24 | 1.92 | 2.44 | 0.92 | 1.98 | 13.6 | 1.99 | 4.58 |
| **4Q97** | 108 | 108 | 39 | 3.42 | 4.88 | 0.64 | 19.77 | 0.0 | 6.71 | 25.11 |
| **4DBH** | 269 | 269 | 26 | 7.92 | 9.64 | 0.75 | 5.51 | 0.0 | 6.13 | 12.03 |
| **2FHE** | 216 | 216 | 84 | 47.46 | 62.22 | 0.99 | 0.51 | 93.8 | 0.58 | 1.1 |
| **1DUG** | 234 | 234 | 35 | 35.11 | 35.87 | 0.98 | 0.83 | 61.3 | 0.83 | 1.93 |
| **3SW1** | 134 | 134 | 57 | 0.0 | 0.0 | 0.57 | 12.93 | 0.0 | 13.37 | 21.55 |
| **3MMH** | 167 | 166 | 26 | 2.5 | 3.09 | 0.57 | 13.97 | 0.0 | 13.47 | 22.6 |
| **4RAZ** | 134 | 134 | 52 | 26.27 | 31.96 | 0.59 | 9.55 | 5 | 7.02 | 21.66 |
| **3GW7** | 215 | 215 | 23 | 0.0 | 0.0 | 0.5 | 24.05 | 0.0 | 24.7 | 55.64 |
| **1VRW** | 289 | 289 | 25 | 2.4 | 2.91 | 0.58 | 12.4 | 0.0 | 12.98 | 31.98 |
| **4EC7** | 108 | 108 | 22 | 0.81 | 0.97 | 0.54 | 12.89 | 2.9 | 12.4 | 23.7 |
| **2C2X** | 280 | 280 | 40 | 6.62 | 8.57 | 0.97 | 1.22 | 22.2 | 1.34 | 2.41 |
| **1VJ2** | 114 | 114 | 27 | 3.85 | 4.63 | 0.98 | 0.65 | 15.2 | 0.67 | 1.32 |
| **4EP4** | 166 | 166 | 27 | 5.47 | 6.48 | 0.56 | 15.9 | 0.0 | 14.81 | 29.29 |
| **1Z3A** | 156 | 156 | 23 | 7.32 | 8.26 | 0.92 | 1.84 | 11.4 | 1.91 | 3.7 |
| **4ZBD** | 219 | 219 | 84 | 20.61 | 24.55 | 0.98 | 1.06 | 42.6 | 0.76 | 2.2 |
| **1PM7** | 199 | 199 | 105 | 30.95 | 45.22 | 0.98 | 0.91 | 55.9 | 0.78 | 1.87 |
| **2JL4** | 212 | 212 | 71 | 30.14 | 36.97 | 0.97 | 1.1 | 65.7 | 1.17 | 2.44 |
| **3NYG** | 93 | 93 | 99 | 40.4 | 51.26 | 0.99 | 0.51 | 69.8 | 0.53 | 1.03 |
| **2HIQ** | 96 | 96 | 29 | 2 | 2.42 | 0.58 | 16.31 | 0 | 11.67 | 22.65 |
| **1Q7G** | 358 | 358 | 43 | 7.64 | 9.52 | 0.97 | 1.51 | 12.5 | 1.61 | 2.97 |
| **1ITU** | 369 | 369 | 27 | 5.48 | 6.3 | 0.74 | 6.33 | 1.9 | 6.15 | 14.41 |
| **3ZJL** | 191 | 191 | 28 | 6.16 | 7.09 | 0.84 | 3.28 | 2 | 3.05 | 7.66 |
| **1DC4** | 323 | 323 | 38 | 5.7 | 6.98 | 0.51 | 22.53 | 0.0 | 20.69 | 56.98 |
| **1SW7** | 245 | 245 | 154 | 42.21 | 65.12 | 0.99 | 0.31 | 88.4 | 0.31 | 0.68 |

Table S6. Detailed results of GD on Set C with predicted inter-chain contacts as input.

| **Target name** | **Length of chain A** | **Length of chain B** | **Number of predicted interchain contacts** | **Precision of predicted interchain contacts (%)** | **Recall of predicted interchain contacts (%)** | **TM-score** | **RMSD** | **f_nat** | **I_RMSD** | **L_RMSD** |
| --- | --- | --- | --- | --- | --- | --- | --- | --- | --- | --- |
| **3KRS** | 249 | 249 | 88 | 37.74 | 45.8 | 0.975 | 1.287 | 56 | 1.325 | 2.726 |
| **1WYI** | 248 | 248 | 141 | 46.28 | 64.93 | 1 | 0.159 | 88.9 | 0.166 | 0.375 |
| **3OGQ** | 112 | 112 | 21 | 5.37 | 5.88 | 0.525 | 14.553 | 0 | 12.173 | 31.829 |
| **1C6X** | 99 | 99 | 151 | 40.24 | 49.28 | 0.994 | 0.403 | 84.6 | 0.4 | 0.915 |
| **2BTM** | 250 | 250 | 121 | 38.5 | 52.17 | 0.998 | 0.392 | 67.3 | 0.433 | 0.825 |
| **4LUL** | 189 | 189 | 60 | 22.84 | 26.62 | 0.981 | 0.975 | 37.8 | 1.026 | 2.47 |
| **2YPI** | 247 | 247 | 107 | 40.34 | 50.71 | 0.995 | 0.569 | 65.9 | 0.595 | 1.174 |
| **3MWS** | 99 | 99 | 140 | 38.95 | 47.86 | 0.994 | 0.392 | 76.8 | 0.375 | 1.021 |
| **2FDE** | 99 | 99 | 157 | 38.73 | 47.52 | 0.993 | 0.435 | 77.8 | 0.418 | 0.896 |
| **3EM6** | 99 | 99 | 149 | 39.31 | 47.89 | 0.995 | 0.39 | 80.4 | 0.376 | 0.913 |
| **3LZU** | 99 | 99 | 157 | 40.46 | 48.61 | 0.991 | 0.492 | 84.5 | 0.466 | 0.94 |
| **3S45** | 99 | 99 | 147 | 45.51 | 52.78 | 0.991 | 0.501 | 84 | 0.439 | 1.014 |
| **4M8Y** | 100 | 100 | 142 | 39.43 | 47.92 | 0.991 | 0.492 | 87.5 | 0.439 | 1.056 |
| **2AOG** | 99 | 99 | 150 | 41.04 | 48.97 | 0.993 | 0.44 | 77.4 | 0.392 | 0.956 |
| **4M8X** | 99 | 99 | 161 | 40.8 | 48.63 | 0.993 | 0.446 | 86.2 | 0.391 | 0.974 |
| **3U7S** | 99 | 99 | 166 | 37.43 | 45.58 | 0.991 | 0.504 | 82.7 | 0.426 | 1.112 |
| **4YMZ** | 250 | 250 | 104 | 40 | 48.65 | 0.988 | 0.876 | 78.9 | 0.881 | 2.044 |
| **2FDD** | 99 | 99 | 160 | 40.11 | 47.65 | 0.99 | 0.538 | 81.8 | 0.492 | 1.27 |
| **4COB** | 206 | 206 | 38 | 5.98 | 7.01 | 0.602 | 16.347 | 0 | 11.312 | 24.4 |
| **3SK2** | 132 | 132 | 44 | 16.2 | 17.68 | 0.587 | 18.453 | 0 | 11.871 | 27.072 |
| **3DSB** | 146 | 146 | 34 | 3.11 | 3.64 | 0.557 | 11.259 | 1.9 | 11.62 | 25.167 |
| **1KPB** | 113 | 113 | 48 | 16.22 | 17.96 | 0.728 | 4.262 | 1.6 | 4.578 | 7.736 |
| **5CPG** | 155 | 155 | 97 | 2.33 | 3.59 | 0.616 | 21.865 | 0 | 15.814 | 29.212 |
| **2E8Q** | 265 | 265 | 81 | 21.74 | 26.32 | 0.959 | 1.705 | 32.8 | 0.729 | 3.307 |
| **1A05** | 357 | 357 | 24 | 10.73 | 11.05 | 0.59 | 11.663 | 4.3 | 9.941 | 25.136 |
| **2E8S** | 265 | 265 | 77 | 21.84 | 25.86 | 0.961 | 1.655 | 34.4 | 0.77 | 3.184 |
| **1LQO** | 134 | 134 | 60 | 15.11 | 17.09 | 0.621 | 7.869 | 0 | 7.883 | 15.129 |
| **1XSE** | 274 | 274 | 70 | 19.21 | 21.67 | 0.985 | 1.02 | 29.3 | 1.06 | 2.13 |
| **1EQU** | 284 | 284 | 21 | 7.58 | 7.77 | 0.963 | 1.639 | 9.1 | 1.603 | 3.361 |
| **3I3G** | 143 | 143 | 69 | 18.03 | 20.39 | 0.54 | 7.959 | 0 | 8.721 | 19.465 |
| **1V5Z** | 217 | 217 | 23 | 7.44 | 7.69 | 0.579 | 14.496 | 0 | 15.796 | 28.149 |
| **3X22** | 217 | 217 | 52 | 9.7 | 11.06 | 0.59 | 12.986 | 3 | 13.98 | 25.484 |
| **3TY2** | 245 | 245 | 57 | 3.41 | 4.17 | 0.584 | 13.647 | 0 | 11.13 | 26.812 |
| **3BM4** | 197 | 197 | 60 | 3.53 | 4.29 | 0.591 | 19.674 | 0 | 7.361 | 28.718 |
| **4R5M** | 369 | 369 | 29 | 5.22 | 5.53 | 0.58 | 11.061 | 4.5 | 8.494 | 25.924 |
| **1MB4** | 369 | 369 | 29 | 5.07 | 5.36 | 0.659 | 10.764 | 2.3 | 6.349 | 18.931 |
| **1QIN** | 176 | 176 | 46 | 12.32 | 12.82 | 0.54 | 11.6 | 0 | 9.482 | 23.708 |
| **4TTB** | 189 | 189 | 33 | 6.46 | 6.79 | 0.591 | 14.135 | 0.9 | 10.907 | 25.925 |
